# Complete and haplotype-resolved maps of genomic and epigenetic discordance in monozygotic twins

**DOI:** 10.1101/2025.10.24.684490

**Authors:** Tengxue Lou, Dongming Fang, Guigao Lin, Zheng Jia, Yafang Qin, Yabin Tian, Lingxin Qiu, Xin Jin, Lei Cheng, Dongya Wu, Kuo Zhang, Chentao Yang, Jie Huang

## Abstract

Telomere-to-telomere (T2T) genome assemblies are indispensable for accurate detection of genetic variation and for resolving complex repetitive regions. Monozygotic (MZ) twin pedigrees provide a powerful model to investigate *de novo* mutations (DNMs), however, comprehensive, haplotype-resolved analyses of structural variation (SV), allele-specific inheritance in complex regions, and DNA methylation in diploid human genomes remain limited. Here, we generated complete, haplotype-resolved T2T assemblies for two female twins (C33 and C35) from a Han Chinese pedigree by integrating complementary, state-of-the-art sequencing technologies. The resulting T2T-C33 and T2T-C35 assemblies are highly contiguous and complete, with Genome Continuity Inspector (GCI) scores of 74.94 (maternal) and 77.94 (paternal), and consensus quality values (QV) >75 (*k* = 21). We comprehensively cataloged 62 inter-twin single-nucleotide variants (SNVs), 15 small indels, and identified both shared and private DNMs, revealing nascent genomic divergence between the MZ twins. Focused interrogation of complex regions uncovered pronounced haplotype-specific length polymorphisms and structural heterogeneity within centromeric higher-order repeat (HOR) arrays. Notably, we observed extensive HOR copy-number variation between haplotypes, including a large copy-number difference on maternal chromosome 18, underscoring dynamic HOR array evolution even among genetically identical individuals. Concurrently, genome-wide DNA methylation profiling delineated allele-specific epigenetic variation that may contribute to phenotypic discordance. Together, these high-quality, diploid T2T genomes from a Han Chinese pedigree provide a valuable resource for population-aware genomics and reveal fine-scale, haplotype-specific divergence in MZ twins. Our results advance understanding of repeat dynamics, centromeric architecture, epigenetic variation and the spectrum of human genomic variation at single-base and structural scales.

## Introduction

The Human Genome Project ushered in the omics era. However, gaps in complex repetitive regions (e.g., centromeres, telomeres, rDNA arrays), which are crucial for genome stability, cell division, and ribosome function and are linked to diseases like chromosomal abnormalities and cancer^1-3^, were present in early references like GRCh37/38. Advances in long-read sequencing (e.g. PacBio high-fidelity (HiFi) and ultra-long Oxford Nanopore Technology (UL-ONT)) and algorithms (e.g. verkko and hifiasm) enabled T2T gap-free genomes^4^, exemplified by the T2T Consortium’s CHM13 haploid assembly, which revealed centromeric HOR structures and rDNA polymorphisms^5^. However, CHM13’s haploid nature limits allelic heterogeneity representation, while HG002’s diploid assembly addressed this but, like GRCh38 and CHM13, is Eurocentric^6^. East Asian (EAS) genomes like CN1 and YAO show significant divergence from CHM13—e.g, ∼2 million undetected rare SNVs in EASs using CHM13 vs. CN1^7^, and YAO’s 326.6/319.7 Mb differences, >3,000 divergent genes, tens of thousands of SVs, and 78 novel genes^8^—impacting precision medicine reliability.

MZs sharing nearly identical genetic backgrounds due to their origin from a single fertilized egg serve as an ideal model for studying genome assembly, somatic variations occurring during individual development, the influence of environmental factors, and resolving the genetic stability of complex genomic regions^9-11^. Recent studies reveal that approximately 15% of MZs exhibit detectable genetic differences arising early in embryonic division, averaging 5.2 mutations, which can contribute to phenotypic discordance such as disease susceptibility^9^. The coordinated changes of such mutations and epigenetic modifications (such as DNA methylation) may be the genetic basis of phenotypic differentiation in twins^12^. Wang et al.^13^ investigated the Chinese T2T-CQ pedigree, comprising an MZ pair and their parents, analyzing the twin offspring data collectively. The team successfully generated a high-quality haplotype-phased T2T genome assembly (QV>70). Through comparative analysis of centromeric sequences between the twins and CHM13, they discovered copy number variations (CNVs) in the HOR arrays among their haplotypes^13^. Notably, the study generated only a merged diploid T2T assembly of the offspring rather than haplotype-resolved assemblies for each twin. This underscores the significant scientific value of utilizing the twin model for separate genome assembly, followed by in-depth analysis of complex genomic regions within twin pairs. However, existing twin genomic studies still find it difficult to accurately capture dynamic events such as centromere recombination, DNMs and allele-specific methylation.

Here, by capitalizing on multi-platform sequencing technologies, we successfully constructed complete T2T diploid genomes for the MZ twin offspring of a Han Chinese family. Leveraging this unique MZ pedigree model, we systematically interrogated inter-haplotype genomic variation, centromeric architecture, allele-specific DNA methylation patterns, and DNMs. Our analyses revealed subtle but significant genetic discordances between the MZ twins at the base-pair and structural levels, when compared to references (CHM13, CN1 and YAO) and population databases (Human Pangenome Reference Consortium (HPRC) and Human Genome Structural Variation Consortium (HGSVC)). While further establishing the pronounced polymorphisms in repeat-rich genomic regions between individuals from distinct populations. Furthermore, allele-specific methylation profiling identified differential methylated regions (DMRs) enriched in regulatory elements and coding sequences. Our work not only delivers a high-quality, T2T diploid reference genome resource for the Han Chinese population but also provides critical theoretical underpinnings for future investigations into the mechanisms of human genomic stability, the dynamic evolutionary patterns of repetitive sequences, and the population-specific genetic and epigenetic architecture of EAS individuals.

## Results

### Diploid genome assembly for the Chinese family

The study constructed a T2T-level diploid genome using a Chinese Han family comprising father (C34), mother (C32) and MZ twin daughters (C33 and C35). Principal Component Analysis (PCA) revealed that the majority of the genomic ancestry of the Han Chinese family clusters with Han Chinese populations when compared to the diverse global populations in the 1000 Genomes Project (1KGP)^14^ (Supplementary Fig. 1). For each individual, multi-platform sequencing strategies were employed, including PacBio HiFi sequencing (115×)^15^, UL-ONT sequencing (80×, including 50× reads >100 kb)^8^ and MGISEQ short-read sequencing (80×, 150 bp paired-end) (Supplementary Table 1).

Existing genome assemblers lack native capacity for T2T assembly. We therefore implemented a hierarchical strategy, initiating with haplotype-resolved assemblies from leading tools (verkko^16^ and hifiasm^17^) using trio-binned datasets. Haplotypes were reconstructed by synthesizing contigs from both assemblers through alignment against CHM13 (Supplementary Fig. 2). Gap resolution was performed using TGS-Gapcloser^18^ with trio-binned UL-ONT reads, the Canu^19^ assembly and a hifiasm assembly generated from the trio-binned ONT reads (Supplementary Fig. 3), reducing gaps to 6 (maternal) and 3 (paternal) for C33, 6 (maternal) and 2 (paternal) for C35, largely confined to centromeric and rDNA regions. Remaining gaps were eliminated by manual reconstruction of local assembly using UL-ONT reads. For both of twin daughters, all 46 chromosomal termini exhibited verified telomeres, with 44 autosomes and the two X chromosomes confirmed gapless and misassembly-free, as well as the mitochondria (Fig. 1a). To improve base-level accuracy, assemblies underwent iterative refinement across five polishing rounds using binned ONT reads, HiFi reads, and MGISEQ short reads (Supplementary Fig. 3). Only homozygous variants consistently identified by both pipelines were incorporated. SVs (≥50 bp) supported by both HiFi and ONT alignments were manually refined. Final diploid genomes of C33 and C35 are 3.03 Gb (maternal) and 3.02 Gb (paternal), comparable to the CQ^13^ and YAO.Mat^8^ reference genome (Table 1). In comparison to CN1, the maternal haplotypes of C33 and C35 were shorter on chromosomes 1, 7, 9, 14-17, and X (Supplementary Table 2), highlighting the genome size variation among Han Chinese individuals.

**Fig. 1.**
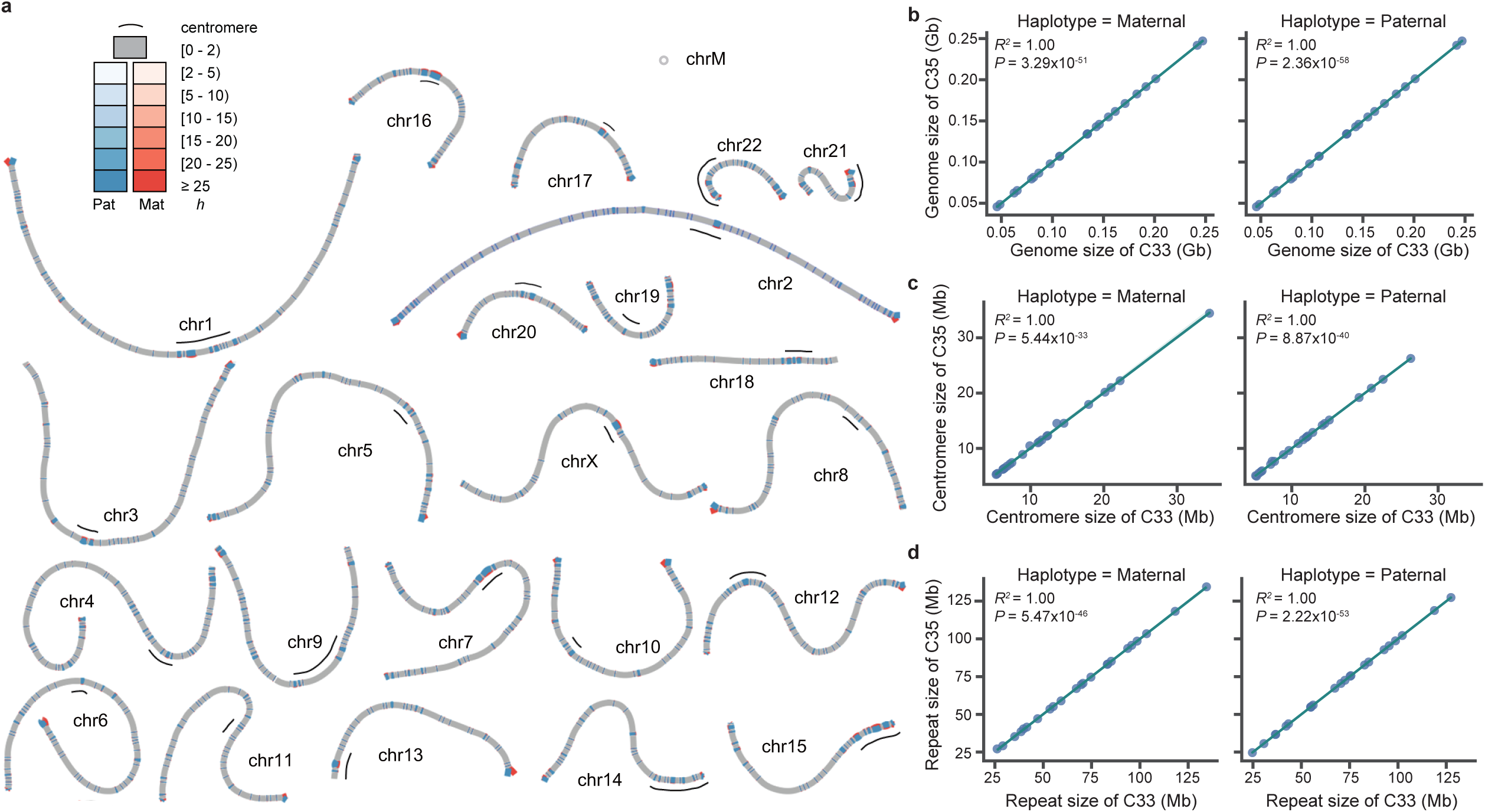
Comparative analysis of genomic characteristics between paternal and maternal haplotypes in C33 and C35. **a** Visualization of heterozygous regions using bubbles, illustrated for the C35 haplotype. Threshold is set at heterozygosity rate (*h* = 2). Homozygous regions are depicted as single grey paths, while heterozygous segments are represented by bubbles according to *h* values, and different *h* values are indicated by varying shades. Notably, centromeres (black lines) exhibit substantially higher *h* values than surrounding regions. **b-d** Scatter plots comparing genome size (b), centromere length (c) and repeat content (d) for maternal and paternal haplotypes between C33 and C35. All comparisons show linear correlation (Pearson correlation coefficients (*R^2^*) = 1.00, two-tailed *P*<1×10^-^³L), indicating near-perfect concordance of haplotype-specific features in the twin pair.

**Table 1.**
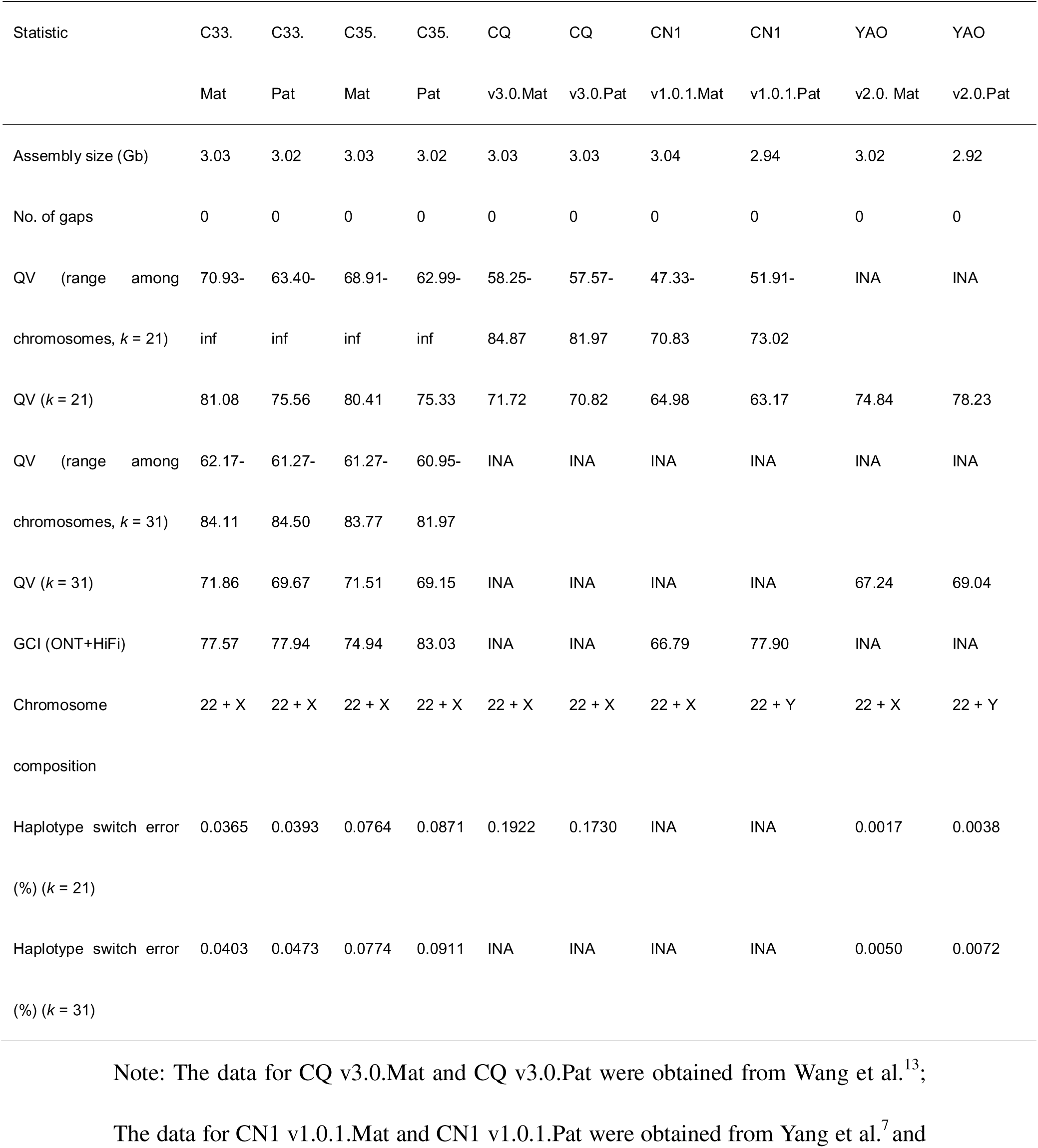
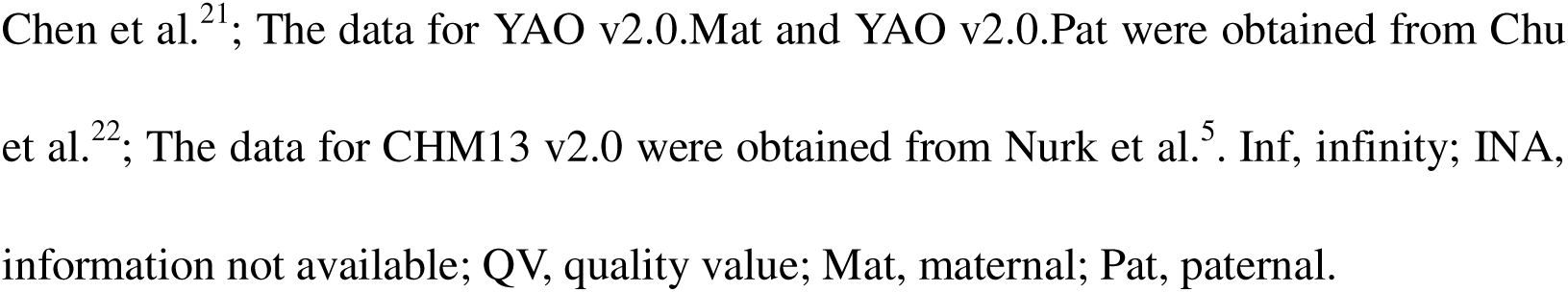
Quality comparison among human telomere-to-telomere genome assemblies.

| Statistic | C33. | C33. | C35. | C35. | CQ | CQ | CN1 | CN1 | YAO | YAO |
| --- | --- | --- | --- | --- | --- | --- | --- | --- | --- | --- |
|  | Mat | Pat | Mat | Pat | v3.0.Mat | v3.0.Pat | v1.0.1.Mat | v1.0.1.Pat | v2.0. Mat | v2.0.Pat |
| Assembly size (Gb) | 3.03 | 3.02 | 3.03 | 3.02 | 3.03 | 3.03 | 3.04 | 2.94 | 3.02 | 2.92 |
| No. of gaps | 0 | 0 | 0 | 0 | 0 | 0 | 0 | 0 | 0 | 0 |
| QV (range among chromosomes, $k = 21$) | 70.93-inf | 63.40-inf | 68.91-inf | 62.99-inf | 58.25-84.87 | 57.57-81.97 | 47.33-70.83 | 51.91-73.02 | INA | INA |
| QV ($k = 21$) | 81.08 | 75.56 | 80.41 | 75.33 | 71.72 | 70.82 | 64.98 | 63.17 | 74.84 | 78.23 |
| QV (range among chromosomes, $k = 31$) | 62.17-84.11 | 61.27-84.50 | 61.27-83.77 | 60.95-81.97 | INA | INA | INA | INA | INA | INA |
| QV ($k = 31$) | 71.86 | 69.67 | 71.51 | 69.15 | INA | INA | INA | INA | 67.24 | 69.04 |
| GCI (ONT+HiFi) | 77.57 | 77.94 | 74.94 | 83.03 | INA | INA | 66.79 | 77.90 | INA | INA |
| Chromosome composition | 22 + X | 22 + X | 22 + X | 22 + X | 22 + X | 22 + X | 22 + X | 22 + Y | 22 + X | 22 + Y |
| Haplotype switch error (%) ($k = 21$) | 0.0365 | 0.0393 | 0.0764 | 0.0871 | 0.1922 | 0.1730 | INA | INA | 0.0017 | 0.0038 |
| Haplotype switch error (%) ($k = 31$) | 0.0403 | 0.0473 | 0.0774 | 0.0911 | INA | INA | INA | INA | 0.0050 | 0.0072 |

Note: The data for CQ v3.0.Mat and CQ v3.0.Pat were obtained from Wang et al.<sup>13</sup>;1046 The data for CN1 v1.0.1.Mat and CN1 v1.0.1.Pat were obtained from Yang et al.<sup>7</sup> and
Chen et al.<sup>21</sup>; The data for YAO v2.0.Mat and YAO v2.0.Pat were obtained from Chu et al.<sup>22</sup>; The data for CHM13 v2.0 were obtained from Nurk et al.<sup>5</sup>. Inf, infinity; INA, information not available; QV, quality value; Mat, maternal; Pat, paternal.

### Genome assessment in MZ twins

Quality assessment using Merqury^20^ and GCI^21^ demonstrated superior accuracy and high continuity in our diploid T2T genomes for twins. For QV (21-mers), the maternal haplotypes exhibit chromosome-wide QV of 80.7 ± 0.4 and paternal haplotypes of 75.4 ± 0.2, exceeding CQ v3.0^13^, CN1 v1.0.1^7^. For QV (31-mers), both maternal haplotypes of C33 (71.86) and C35 (71.51) were higher than YAO v2.0.Mat (67.24) by 5 QV units, whereas the paternal haplotypes of C33 (69.67) and C35 (69.15) were similar to YAO v2.0.Pat (69.04) and YAO v2.0^22^ (Table 1). Notably, one-third of chromosomes yield infinite QV (zero *k*-mer errors), and switch-error rates are 15-30% lower than CQ v3.0 (Supplementary Table 3, Table 1, Supplementary Fig. 4). Despite multiple rounds of polishing, a small number of residual base-level errors persist. *K*-mer spectrum-based analysis of the polished assemblies indicates a marked regional bias. Residual errors are strongly regional and predominantly enriched in centromeres (C33.Mat: 31.84%; C33.Pat: 83.59%; C35.Mat: 44.44%; C35.Pat: 92.71%) and subtelomeres (C33.Mat: 57.40%; C33.Pat: 11.75%; C35.Mat: 25.00%; C35.Pat: 5.68%) across maternal and paternal haplotypes in twins (Supplementary Figs. 5-11), indicating that current polishing algorithms remain sub-optimal for highly repetitive DNA. To further evaluate the assembly continuity of twins, we employed GCI^21^. Our C33 and C35 assemblies achieved GCI scores (ONT+HiFi) of 77.57 (C33.Mat), 77.94 (C33.Pat), 74.94 (C35.Mat) and 83.03 (C35.Pat), surpassing CN1’s values of 66.79 (Mat) and 77.90 (Pat) (Table 1). These results demonstrate the high accuracy, continuity, and completeness of the C33 and C35 genome assemblies.

UL-ONT reads (>100 kb) provided uniform depth across chromosomal (Supplementary Figs. 12-15). To benchmark assembly completeness in clinically refractory loci, we examined 195 genes previously shown to have <90% short-read coverage^23^. Alignment to the T2T reference revealed full-length, single-haplotype representation of segmentally duplicated or highly homologous sequences, including Mucin 1 (*MUC1*), survival motor neuron gene 1 (*SMN1*) and zonadhesin (*ZAN*) (Supplementary Fig. 16). ONT coverage across these regions was devoid of dropouts or allelic collapse, confirming that the C33/C35 diploid assemblies accurately resolve complex medically relevant sequences.

The construction of a reference genome from twin samples has natural genetic consistency. Further comparative analysis of genomic characteristics between paternal and maternal haplotypes of twins revealed perfect correlation in both genome size (maternal: Pearson correlation coefficient *R^2^* = 1.00, *P* = 3.29×10^-51^; paternal: *R^2^* = 1.00, *P* = 2.36×10^-58^) and centromere size (maternal: *R^2^* = 1.00, *P* = 5.44×10^-33^; paternal: *R^2^* = 1.00, *P* = 8.87×10^-40^), with significant correlations in non-repeat sizes (maternal: *R^2^* = 1.00, *P* = 5.47×10^-46^; paternal: *R^2^* = 1.00, *P* = 2.22×10^-53^) (Figs. 1b-d), confirming the expected high genomic concordance.

### Genomic comparison with published human T2T references

To investigate structural variants between twins and the previously released T2T genome assemblies of Chinese Han individuals (CN1 and YAO), we compared variants, such as SNVs, indels, inversions and translocations, across haplotypes. This analysis revealed that the maternal and paternal haplotypes of C33 and C35 shared more identical variants than with YAO or CN1, while also harboring distinct variants unique to each haplotype (Fig. 2a). Further, collinearity analysis revealed that these differences were primarily concentrated in centromeric regions, with proximal centromeric segments on several chromosomes harboring unalignable sequences, suggesting assembly issues in these areas (Supplementary Fig. 17). Subsequent comparative analysis, masking both centromeric and telomeric sequences, showed a 60% reduction in the genome-wide variant count (Fig. 2b), further indicating that the majority of the observed differences reside within centromeric and subtelomeric regions. The results indicated that beyond the variants shared among all haplotypes, both maternal and paternal haplotypes of twins C33 and C35 shared more variants than with YAO or CN1, while each haplotype also possessed unique variants (Fig. 2b), revealing the genetic variation present among Asian individuals. Given that individual SVs are more likely than individual SNVs or indels to affect gene function^24^. Inversion variants, as significant SVs, can markedly influence gene function and phenotype by altering gene structure or regulation, and modulating gene expression^25^. We performed alignment against YAO to identify all inverted regions exceeding 50 kb (Supplementary Tables 4-5). Frequency distribution analysis using the HPRC^26^ and HGSVC^27^ databases, combined with T2T genomes (CHM13, HG002, CN1 and YAO), showed that the consistency rates of these large inversions with the maternal haplotype of twins ranged from 10.27% to 99.72%, while consistency with the paternal haplotype ranged from 17.04% to 100% (Supplementary Tables 4-5). This variability reflects the polymorphic nature of these genomic regions.

**Fig. 2.**
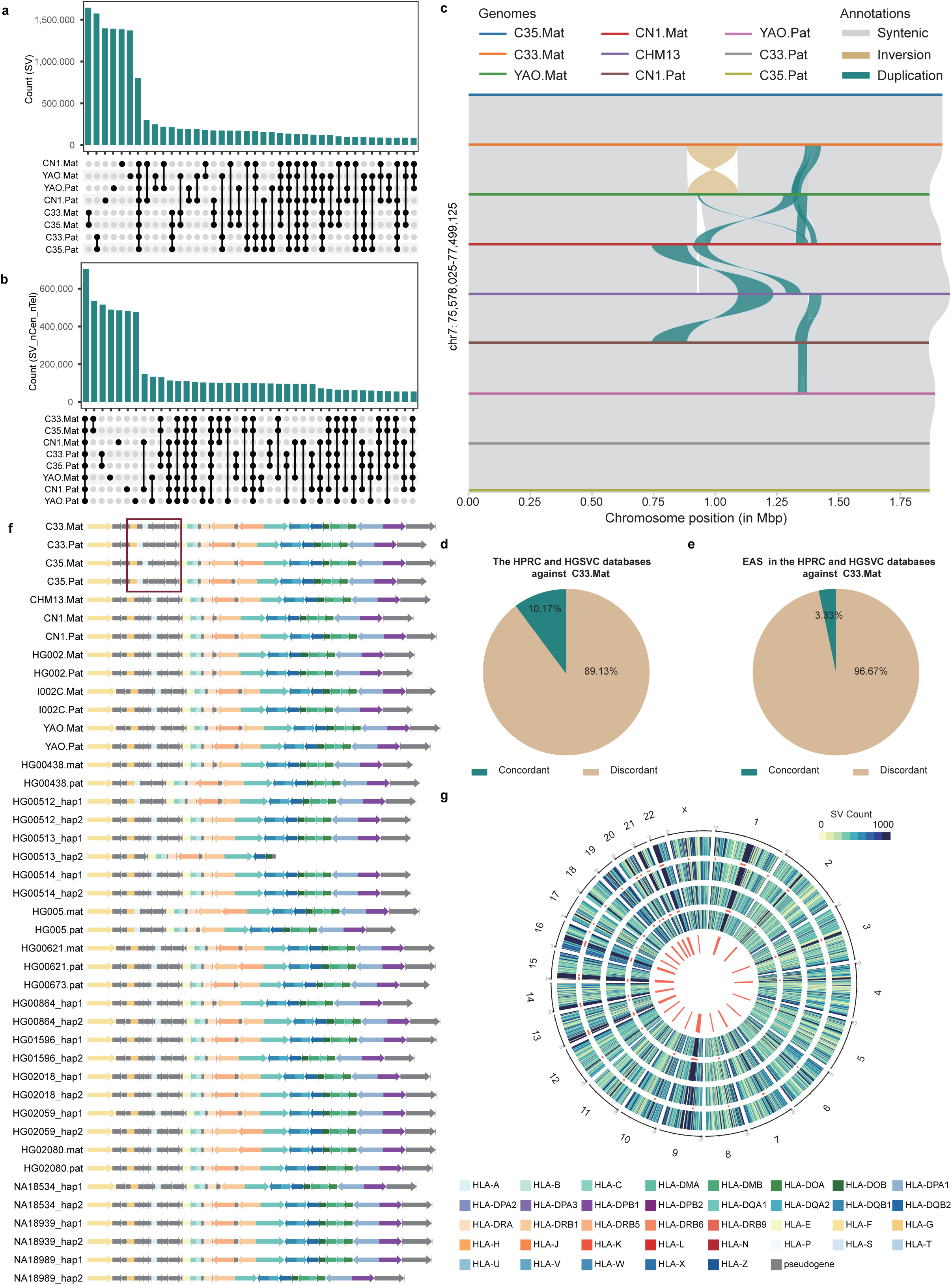
Comparative genomic analysis of variants between twins and population. **a** Genome-wide variant analysis (SNVs, indels, inversions, translocations) across eight haplotypes (C33, C35, YAO, and CN1) including centromeric regions. **b** Genome-wide variant analysis across eight haplotypes, with centromeres excluded. bar plot quantifies variants shared among haplotypes. **c** Twin-specific structural variant highlighting a 1.8-Mb chromosomal segment, featuring an ∼800 kb inversion (brown) unique to both C33.Mat and C35.Mat haplotypes, with syntenic alignments to YAO, CN1, and CHM13. **d** Frequency of a chr7 inversion in C33/C35.Mat haplotype compared to global haplotypes in HPRC and HGSVC databases. **e** Concordance of the same inversion in C33/c35.Mat within East Asian (EAS) populations from HPRC/HGSVC. **f** Allele-specific comparison of the HLA region in twins versus reference individuals or cell line. Each row represents an haplotype. Alleles are depicted as colored arrows corresponding to specific gene families. The red box denotes a genomic segment exhibiting allelic discordance between the twins in comparison to the population. **g** Circos plot displaying variant distribution. From outer to inner, the concentric layers represent the CHM13 reference chromosomes, followed by the distribution of variants in C33.Mat, C35.Mat, C33.Pat, and C35.Pat haplotypes, concluding with the centromeres (red line). Chi-square tests (*P*<0.01) indicate significant divergence between twin haplotypes at centromeric and subtelomeric regions (red dots).

Notably, comparative analysis revealed 1.8-Mb inversions specifically present in the maternal haplotypes of C33 and C35, which were absent in the YAO, CN1, and CHM13 haplotypes as well as in the paternal haplotypes of twins (Fig. 2c). Careful validation of the assemblies for both the maternal and paternal haplotypes of the twins provides strong support for these inversions, as evidenced by ONT read coverage (Supplementary Fig. 18). Furthermore, comparison of the chromosome 7 inversion in the C33.Mat haplotype with the HPRC^26^ and HGSVC^27^ databases showed a low concordance rate of 10.17%. Strikingly, concordance with EAS individuals within these databases was merely 3.33% (Figs. 2d-e, Supplementary Tables 6-7), indicating that the inversion pattern in C33.Mat diverges not only from global genomic backgrounds but also from EAS-specific variations. Human leukocyte antigens (HLAs), encoded by a highly polymorphic gene cluster on chromosome 6, are essential for adaptive immunity^28^. To perform a high-resolution haplotype-level comparison of HLA gene variation, we compared the genotypes between twins, EAS individuals from HPRC^26^ and HGSVC^27^, European individual HG002 and European cell line CHM1<u>3. Th</u>e results revealed high allelic concordance across most HLA genes. However, a specific region of noteworthy discordance (highlighted in red, Fig. 2f) revealed divergent gene organization, including variations in gene order or copy number, between the twins and other haplotypes. The validation of the maternal and paternal haplotype assemblies confirmed the accuracy of the HLA regions, which was demonstrated by the ONT read coverage (Supplementary Fig. 19). Further analysis indicated that this region is enriched with pseudogenes and repetitive sequences (Fig. 2f, Supplementary Fig. 20), a phenomenon likely attributable to extreme local polymorphism. Additionally, homologous variant analysis identified differential SNVs, indels, and SVs between twins, predominantly enriched in centromeric regions (Fig. 2g). These findings underscore inter-individual genomic divergence, even within MZ twins.

### Centromeric annotations and unique HOR patterns in twin haplotypes

The centromere and pericentromeric regions are dominated by long, highly homologous tandem repeat sequences (satellite DNA). To minimize their confounding effects on downstream analyses, we first annotated alpha-satellite arrays in C33 and C35 and characterized HORs on all non-acrocentric chromosomes. The annotated centromeric spans ranged from 5.06 to 26.25 Mb (paternal) and 5.28 to 34.44 Mb (maternal) in C33, and 4.97 to 26.25 Mb (paternal) and 5.28 to 34.43 Mb (maternal) in C35 (Supplementary Table 8). High-resolution sequence-identity plots generated with StainedGlass^29^ revealed organizational differences in tandem repeat organization between the twin haplotypes (Supplementary Fig. 21).

Alpha-satellite HOR array lengths are known to vary across human population ^1,30^. To quantify array length variation in C33 and C35 and to compare these with the CHM13 reference, we analyzed the HOR array lengths for each haplotype. In C33, HOR lengths spanned 1.69-6.93 Mb (maternal) and 1.72-9.13 Mb (paternal). While in C35, they ranged 1.69-7.21 Mb (maternal) and 2.41-9.13 Mb (paternal). By contrast, CHM13 HOR lengths range 2.11-8.38 Mb (Supplementary Table 9). Comparative analysis revealed length differences between each twin haplotype and CHM13, with paternal and maternal haplotypes also exhibiting length differences across all chromosomes except for chromosomes 1, 7 and 12 (Supplementary Table 9, Fig. 3a). These results indicate extensive HOR length polymorphism both between the twin haplotypes and relative to CHM13.

**Fig. 3.**
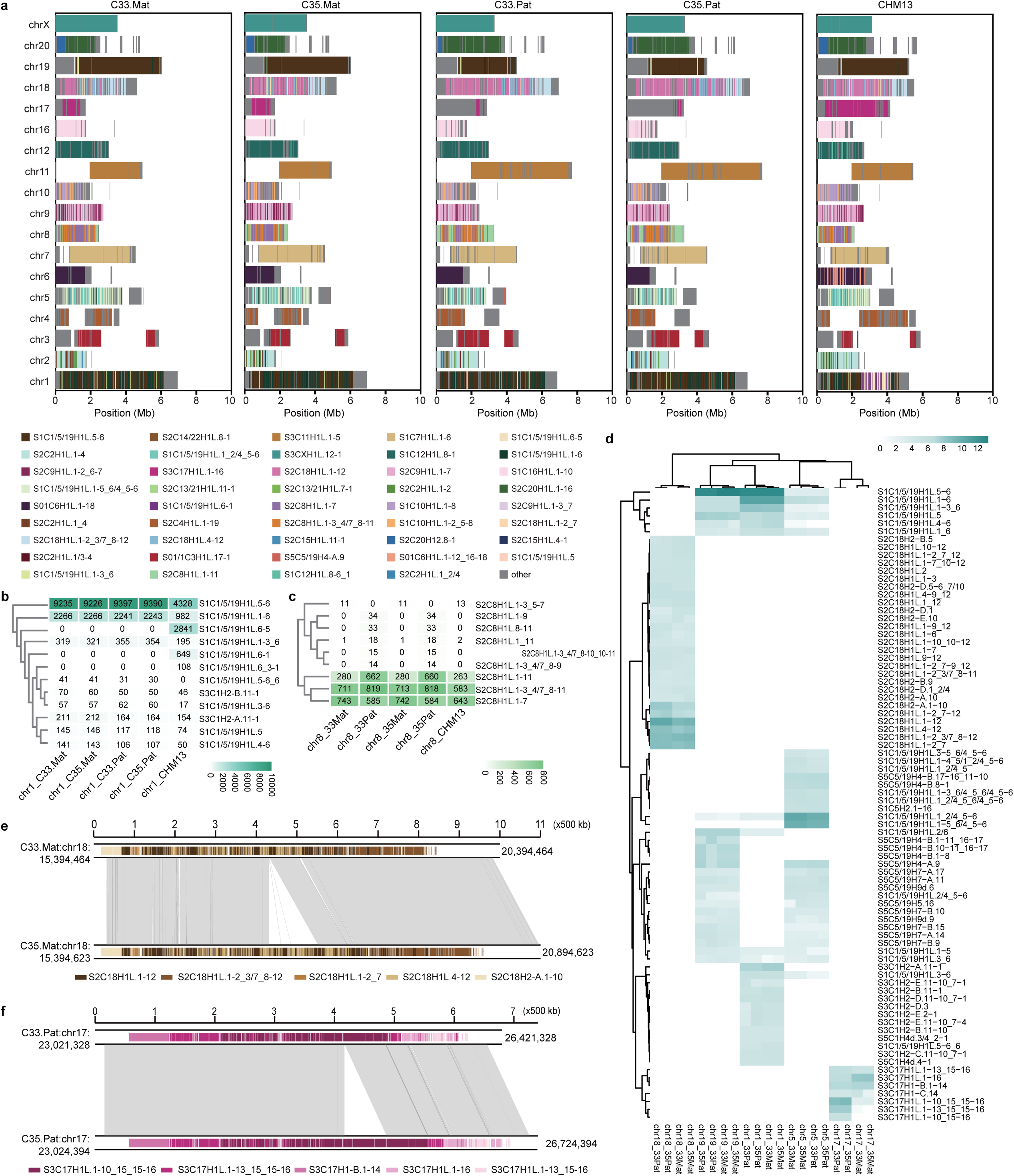
Comparative analysis of higher-order repeat (HOR) array structure in centromeres across the CHM13 reference genome and twin haplotypes. **a** The HOR array structure is shown on the axes, along with the organization of each centromeric region in C33.Mat, C33.Pat, C35.Mat, C35.Pat and CHM13. **b, c** Heatmaps comparing the HOR number on chr1 (b) and chr8 (c) across CHM13, C33.Mat, and C33.Pat. **d** A comparative heatmap displaying chromosomes with HOR differences between C33 and C35. **e, f** Local synteny of centromeres on maternal chr18 (e) and paternal chr17 (f) between twins, revealing copy number variations between haplotypes.

We also observed marked SVs in HOR organization between the twin genomes and the CHM13 reference genome (Fig. 3a). For example, chromosome 1 of CHM13 contains a block of unique HOR sequence composed primarily of specific 2-mer (S1C1/5/19H1L.6-5, S1C1/5/19H1L.6-1) and 3-mer (S1C1/5/19H1L.6_3-1) units that are absent from both maternal and parental haplotypes of the twins (Fig. 3b). Conversely, chromosome 8 of the twins contains five unique paternal-haplotype-specific HOR units (S2C8H1L.1-9, S2C8H1L.8-11, S2C8H1L.1_11, S2C8H1L.1-3_4/7_8-10_10-11, and S2C8H1L.1-3_4/7_8-9) that are not present in the twin maternal haplotype or in CHM13 (Fig. 3c). Taken together, these observations demonstrate substantial HOR structural polymorphism across multiple loci, manifested as haplotype-specific absence or CNV of distinct HOR units.

Analysis of centromeric SVs within twin homologous haplotypes revealed that all chromosomes exhibited variable HOR copy number differences (Fig. 3d), with particularly pronounced variation on chromosomes 17 and 18 (Figs. 3e-f). Assembly validation of both haplotypes of the twins, supported by ONT read coverage, corroborates these structural differences (Supplementary Fig. 22). Local collinearity analysis identified 402 CNVs (*P*<0.05) between the maternal haplotypes of C33 and C35 on chromosome 18, predominantly localized to the S2C18H1L.1-12, S2C18H1L.1-2_3/7_8-12, S2C18H1L.1-2_7, and S2C18H2-A.1-10 units. By contrast, the paternal haplotypes on chromosome 17 predominantly harbored a 13-mer HOR (S3C17H1L.1-10_15_15-16), with 152 CNVs observed between twins (*P*<0.05) (Figs. 3e-f, Supplementary Table 10). Notably, the major component of chromosome 17 in the CN1 genome, a 13-mer HOR (S3C17H1L.15#), was undetected^7^. These findings reveal HOR copy number polymorphism in twin centromeric regions and distinct patterns of variation between maternal and paternal haplotypes.

### Allele-specific DNA methylation in MZ twins

Studies have shown that phenotypic differences between MZ twins are driven not only by genetic variation but also by epigenetic processes, notably DNA methylation^31^. DNA methylation plays an important role in biological processes in human health and disease^32^. Analysis of cytosine guanine dinucleotide (CpG) count across parental haplotypes in MZ twins C33 and C35 dispalyed differential distribution on all autosomes and chromosome X, with minimal variation between co-twin haplotypes (Supplementary Fig. 23). Chromosome-scale circular plots further indicated broadly similar methylation patterns between twin homologous haplotypes (Fig. 4a). Furthermore, at the genome scale, CpG methylation exhibited a bimodal distribution, with notable disparities in methylation levels observed between homologous haplotypes (Fig. 4b). A notable parent-of-origin-specific expression pattern was observed, with C33 exhibiting high expression on the maternal haplotype and C35 showing preferential expression on the paternal haplotype, likely due to parental-origin effects linked to X-chromosome inactivation^32^.

**Fig. 4.**
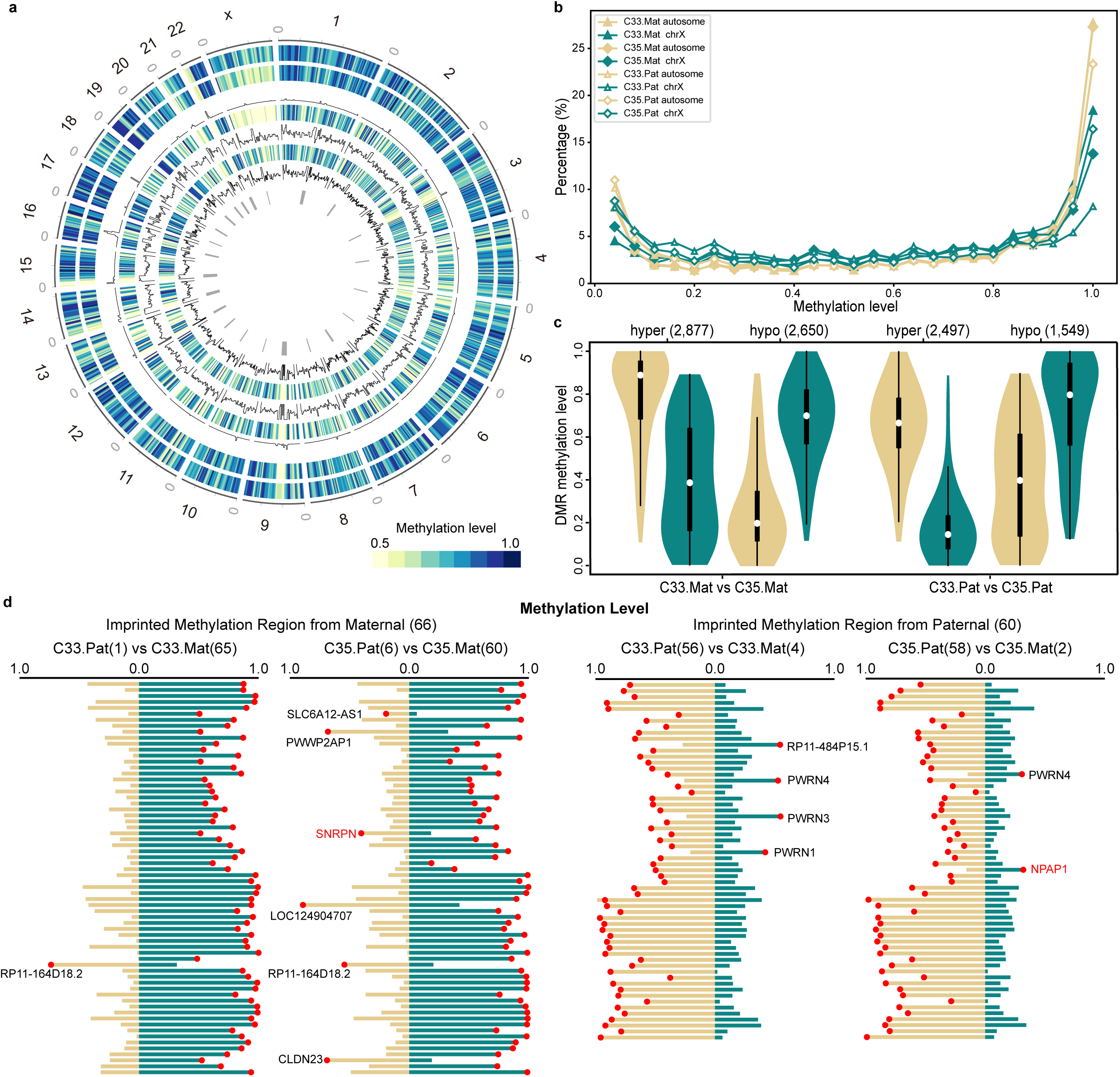
Methylation analysis between the twins. **a** Circular visualization of chromosomed-scale methylation levels aligned to the C33.Mat genome. From outer to inner, the concentric layers represent methylation profiles and single-nucleotide variant (SNV) density plots for both maternal and paternal haplotypes of C33 and C35, as well as centromere positions (dark gray bars). Methylation levels are depicted using a color gradient, while SNV densities are shown as vertical ticks. **b** Compare genome-wide CpG methylation levels between the monozygotic twins for each haplotype. **c** Comparison of methylation levels in differentially methylated regions (DMRs) between maternal and paternal haplotypes in twins. The white dot inside each violin represents the median value, and the thick black bar represents the interquartile range. The number of DMRs in each category is indicated in parentheses. **d** Validation of parent-of-origin (PofO)-specific DNA methylation patterns in blood-derived imprint control regions. Each horizontal black bar represents a single CpG site, with its length corresponding to genomic position. Red dots denote methylation levels, with the y-axis scaled from 0.0 (unmethylated) to 1.0 (fully methylated). Annotated genes inconsistent with previous study^77^ are highlighted in red.

Analysis of genome-wide differentially methylated cytosine (DMC) distribution patterns revealed that distinct epigenetic discordance between homologous haplotypes across multiple chromosomes in twins (Supplementary Fig. 24a). Furthermore, both maternal and paternal DMCs were enriched in regulatory elements, particularly upstream of transcription start sites (TSS) and coding exons (Supplementary Figs. 24b, c). Further, we identified DMRs: in the maternal comparison (C33 maternal versus C35 maternal) there were 2,877 hypermethylated and 2,650 hypomethylated regions, and in the paternal comparison there were 2,497 hypermethylated and 1,549 hypomethylated regions (Fig. 4c, Supplementary tables 11-12). Unlike DMC distribution patterns, DMRs were significantly enriched in coding exons and untranslated regions (UTRs) relative to TSS upstream regions (Supplementary Fig. 25). This indicates that parental-specific DNA methylation changes have distinct genomic targets, potentially influencing gene expression through different mechanisms. Kyoto Encyclopedia of Genes and Genomes (KEGG) pathway enrichment of DMR-associated genes highlighted overrepresentation of pathways including cancer (Supplementary Fig. 26). Together, these results indicate a largely uniform CpG distribution but measurable allele-specific methylation differences between haplotypes.

We next investigated the methylation pattern differences in centromere regions among the alleles. Consistent with findings in CHM13, we observed a reduction in DNA methylation within the active HOR subregions compared to their flanking sequences across all chromosomes (Supplementary Figs. 27-28), termed as centromeric dip region (CDR). Generally, CDRs typically co-localize with Centromere Protein (CENP) binding sites, consequently linking them to kinetochore attachment^33^. Differential methylation levels of CDR between twins were observed on chromosomes 17 and 21, which also differed from the expression patterns of CHM13^5^ and CN1^7^, probably due to the differences in the major components in the active HOR^7^. In addition, the overall methylation level of the CpG sites in the mitochondrial DNA (mtDNA) was lower than that of nuclear DNA (nDNA), and there was no differentially methylated CpG positions were detected between twins (*P*>0.05 in chi-square test; Supplementary Fig. 29).

Moreover, to characterize allele-specific methylation (ASM) and genomic imprinting patterns, we profiled the methylation level of 66 and 60 imprinted methylation regions from maternal and paternal haplotype, respectively^34^. Analysis of these methylation regions in twins revealed no erroneous switch events. In keeping with canonical imprinting, the active allele was mostly consistently hypomethylated while the silent allele was hypermethylated, confirming methylation-dependent monoallelic repression, such as *NAA60* (Fig. 4d, Supplementary Fig. 30), indicating that the imprinting control regions in blood samples possess stable parent-of-origin-dependent methylation patterns. While this signature was absent for a subset of genes (Fig. 4d, Supplementary Tables 13-14), underscoring tissue- or context-specific imprinting dynamics that may reflect parent-of-origin switching.

### DNMs in MZ twins with associated CpG methylation changes

Recent high-throughput sequencing studies utilizing the “trio” (father-mother-offspring) design have begun to unravel the origin and properties of DNMs, which are increasingly recognized as important contributors to disorders such as neurodevelopmental disorders and congenital heart disease^9,35-36^. DNMs are defined as genetic mutation present in the offspring but absent in both parents. Human DNMs encompass germline and postzygotic mutation (PZMs), which can be catalogued as those shared between twins or private to an individual (Fig. 5a). To identify DNMs in the twin pairs, we applied a stringent, trio-based detection pipeline adapted from established methods^37^. Analysis excluding centromeric regions revealed that C33 and C35 shared 50 SNVs and 8 indels, corresponding to a genome-wide DNM rate of 1.92×10^-^L per site. C33 carried 5 private SNVs and 3 indels (rate: 2.65×10^-^L), whereas C35 harbored 7 private SNVs and 4 private indels (rate: 3.64×10^-^L) (Fig. 5b). This indicated that the shared mutations likely originated from early embryonic events common to both samples, while the private mutations reflect later, lineage-specific accumulation.

**Fig. 5.**
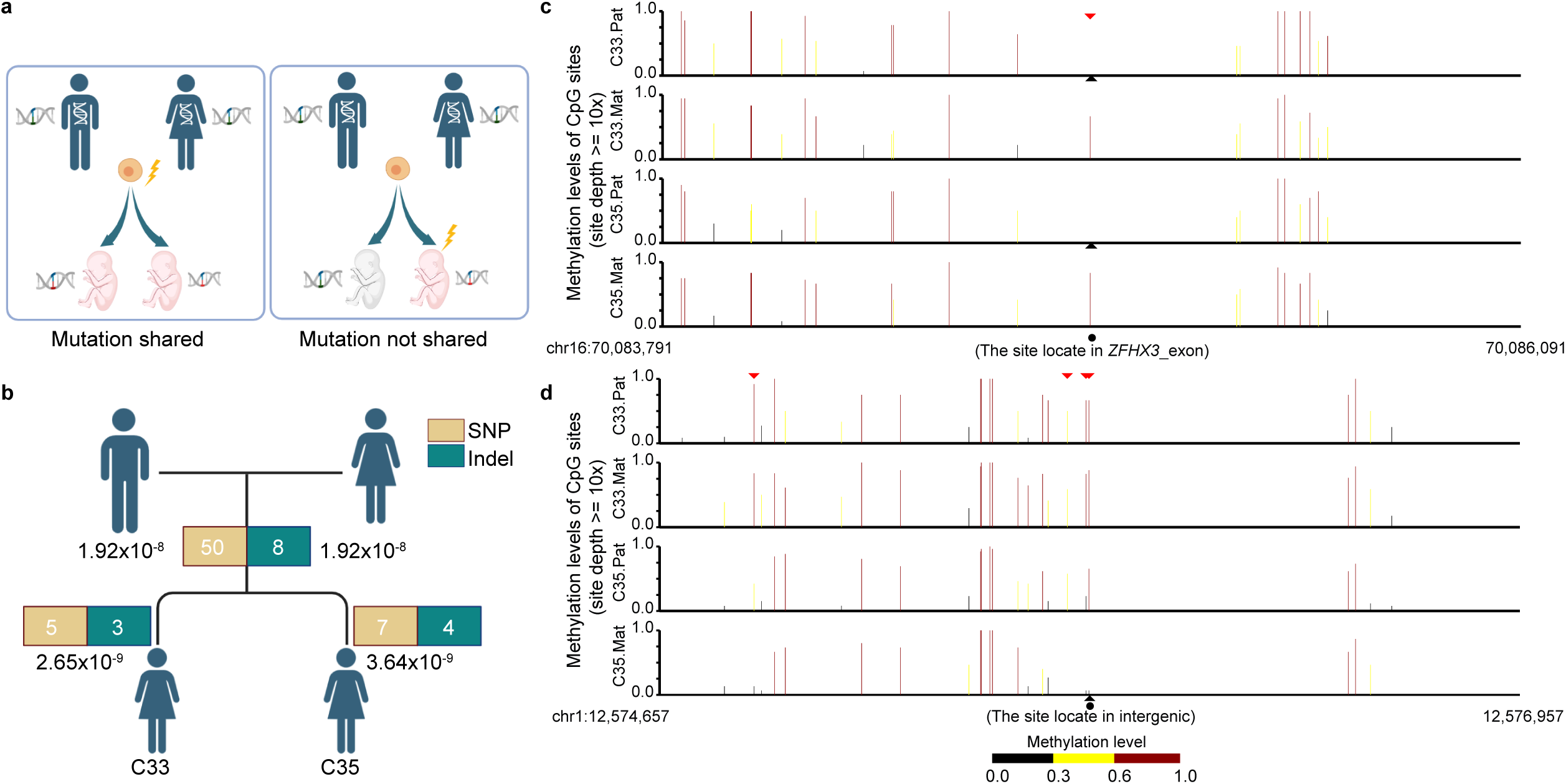
Analysis of *de novo* mutation types in C33 and 35. **a** Schematic representation of the two classes of *de novo* mutations identified in the twin pair: shared mutations (left) and non-shared mutations (right). **b** Counts of *de novo* SNVs and indels detected in each twin. Observed counts are indicated above bars, with mutation rates per site shown below. **c** Methylation profile of *ZFHX3* alleles, showing a *de novo* G>C mutation at a CpG site within an exon of *ZFHX3* (chr16). **d** Methylation profile of an intergenic region (chr12), showing a *de-novo* G>A mutation at a CpG site. Bar plots show the methylation levels for the paternal and maternal haplotypes in each twin. The position of mutant base is marked with a black circle. Black triangles indicate haplotype-specific mutation sites. Red triangles denote CpG sites with allele-specific methylation differences.

Notably, a shared *de novo* G>C substitution was identified within an exon of *ZFHX3* (Zinc Finger Homeobox 3) on chromosome 16 in the paternal haplotype of twins, which was associated with a locus-specific loss of methylation (Fig. 5c). *ZFHX3* encodes a transcription factor involved in cell differentiation and tumor suppression^38^. Previous studies have shown that DNA methylation of *ZFHX3* plays a critical role in the pathogenesis of various diseases, including myocardial infarction, ischemic stroke and pancreatic cancer^39-41^. Consequently, methylation loss at this site may disrupt normal gene regulation and contribute to phenotypic variation. Additionally, a *de novo* G>A mutation was also detected specifically on the maternal haplotype of C35, located within an intergenic region on chromosome 1 and similarly correlated with a loss of DNA methylation (Fig. 5d). Although intergenic, such methylation changes can influence gene regulation by altering chromatin accessibility, modulating enhancer or silencer activity, or disrupting long-range chromosomal interactions, thereby potentially affecting expression of neighboring genes^42-43^. Sequence visualizations confirmed the offspring-specific *de novo* SNVs in *ZFHX3* (Supplementary Fig. 31) and the intergenic region on chromosome 1 in C35 (Supplementary Fig. 32). These findings suggested that DNMs occurring in distinct genomic contexts, both exonic and intergenic, may lead to localized epigenetic dysregulation, highlighting a potential mechanism by which spontaneous genetic variation may influence phenotypic outcomes through altered DNA methylation patterns.

## Discussion

The selection of MZ twins as the foundational material for a complete genome assembly and subsequent epigenomic analysis provides a uniquely high-fidelity biological control. MZ twins are genetically near-identical, sharing essentially 100% of their inherited sequence, which fundamentally enhances the credibility of DNM discovery and allows researchers to reliably attribute differences in phenotype or epigenetic status to postzygotic, environmental, or stochastic factors^10,44^. This methodology stands in contrast to comparisons between unrelated individuals or reliance on haploid-like reference genomes (e.g., T2T-CHM13), significantly strengthening the mechanistic inferences drawn from the observed divergence. By leveraging an innovative Han MZ family model, this study substantially improved diploid genome assembly quality: the twins’ high inter-haplotype homology (>99.9%) provided an intrinsic mutual calibration standard, and direct comparison of twin haplotypes allowed correction of small variant sites that reduced haplotype switch error rates to 0.04% (C33) and 0.08% (C35) (for 21-mers) (Table 1), which was a 4.75-fold improvement over CQ v3.0 (0.19%)^13^. However, the pursuit of the gold-standard QV100 — representing virtually zero errors — remains elusive, underscoring fundamental technical limitations. Our analysis confirms that the centromeric α-satellite DNA and HOR arrays constitute the primary barrier to absolute accuracy^1,45^. These regions are defined by massive, near-perfect repetitive units (often differing by less than 5% from consensus). *K-*mer spectrum analysis revealed a pronounced regional bias in error distribution, with 60% of erroneous *k-*mers concentrated in centromeric regions, 27% in subtelomeric regions, and only 13% elsewhere (Supplementary Fig. 5), consistent with repeat-associated assembly challenges reported by previous studies^1,22,46^ and indicating persistent difficulties in resolving higher-order repeats. The combined effect of this extreme repetitiveness and residual sequencing error rates presents an intrinsic computational challenge in distinguishing true biological variation from assembly artifacts, confirming that centromeres remain the final frontier in achieving perfect linear reference quality. To address these limitations, we recommend integrating microfluidic chromosome-sorting approaches that use dielectrophoresis for single-chromosome resolution^47^, long-read, long-range sequencing technologies (e.g., Hi-C/Pore-C) to span complex loci^16^, and repeat-aware assembly algorithms, such as graph-phasing methods that exploit CENP-B binding sites to delineate α-satellite monomer boundaries^30^.

The utility of a reference genome is fundamentally dependent on its ability to represent global human diversity. Current high-quality resources, such as the T2T-CHM13 haploid assembly and the commonly used variant baseline HG002, primarily reflect European ancestry, creating significant “reference bias” that diminishes the accuracy of variant calling and clinical interpretation in other populations^7^. While there are several human T2T genomes for Asian population^7,13,22^, the high-quality genomes are still insufficient for such a large population. Our complete, haplotype-resolved T2T assemblies from a Han Chinese pedigree represent supplementary diversity in terms of clinically refractory loci. The identification of SVs, such as a 1.8-Mb inversion in the maternal haplotypes of C33/C35 and divergent HLA organization between twins and other EAS haplotypes (Figs. 2c, e, f), underscores notable structural-variation diversity within the Han population. Practically, they highlight the value of ancestry-matched reference genomes in precision medicine: using a phylogenetically proximal reference improves rare-variant detection accuracy by 17.3% and reduces annotation errors for clinically relevant variants by 32.6%^48^. The T2T resolution achieved here is essential for accurately capturing these population-specific SVs and complex repeat elements, which are systematically missed by short-read approaches mapped to incomplete linear references. Furthermore, this high-quality, phased data is indispensable for the advancement of Artificial Intelligence (AI) and Machine Learning (ML) in genomics. AI models, which are increasingly vital for discerning intricate patterns in genomic data, especially SV, require standardized, high-fidelity datasets as "ground truth" for training^49-50^. Training AI models on T2T assemblies significantly improves the identification and interpretation of SVs across diverse cohorts by reducing false positives and negatives inherent to older assemblies^27^. The diploid and population-aware nature of T2T-C33/C35 provides the structural context needed to overcome algorithmic bias and ensure that AI-driven clinical tools are relevant and equitable for diverse global populations.

The traditional definition of MZ twins as genetically identical is substantially refined by the high-resolution power of T2T sequencing. Our focused interrogation of complex regions revealed nascent genomic divergence that challenges previous assumptions of stability^51^. Specifically, the centromeric HOR arrays, essential for accurate chromosome segregation, demonstrated pronounced haplotype-specific length polymorphisms and extensive CNV, including a large CNV difference observed on the maternal chromosome 18 (Fig. 3e). The discovery of such structural dynamics in genetically near-identical individuals underscores that these regions are highly unstable even within the short developmental timescale following zygote splitting. This instability suggests the existence of mechanisms for rapid postzygotic evolution or differential selective pressures acting on centromere structure in distinct cell lineages. Additionally, we identified 50 shared SNVs and 8 shared indels between the twins, alongside twin-specific variants (C33 harboring 5 private SNVs/3 indels; C35 carrying 7 private SNVs/4 indels) (Fig. 5b). These observations align with prior reports in an MZ cohort from Feng et al.^36^, where both shared and private DNMs were found in a schizophrenia twin. Beyond nascent genetic and structural divergence, our study provides the first comprehensive, genome-wide view of the epigenetic landscape on complete, haplotype-resolved assemblies, revealing extensive ASM that likely contributes to phenotypic discordance between MZ twins. This extends previous evidence that epigenetic differences accumulate with age in MZ twins and correlate with divergent susceptibility to complex diseases^52^. Notably, variants in *ZFHX3* co-localized with locus-specific methylation changes (Fig. 5c). This pattern accords with Jonsson et al.^9^, who reported genomic disparities between MZs based on whole-genome sequencing, including differences in somatic mutations and epigenetic state. By anchoring the ASM to haplotype-resolved assemblies, we delineate a mechanistic framework whereby subtle *de novo* genetic variations or structural heterogeneity on one haplotype predispose that allele to differential epigenetic modification, thereby providing a plausible basis for non-heritable phenotypic differences.

These Han Chinese MZ twin assemblies constitute a high-fidelity resource for human genomics, pangenome construction, and the implementation of personalized medicine. Their value is especially acute given that the current gold-standard SV benchmarks, such as those derived from the HG002 genome, overwhelmingly represent European ancestry, leaving a critical gap in EAS specific benchmarks, and constraining clinical interpretation and diagnosis of diseases in these populations. The T2T-C33/C35 datasets, together with associated cell lines, provide the necessary high-fidelity baseline for establishing a comprehensive genetic variation profile for EAS populations. They enable construction of representative pangenome graphs that faithfully model complex, population-specific genetic architectures in Han Chinese individuals, overcoming limitations of European-centric linear reference. By providing a complete, fully phased view of previously intractable regions, this resource supports precise investigations of allele-specific effects in a diploid context. Ultimately, the high-resolution catalogs of DNMs, dynamic structural changes, and allele-specific epigenetic marks advance fundamental understanding of human evolution, while establishing the foundation for population-aware diagnostics and equitable, personalized therapeutic strategies globally.

## Methods

### Sample and immortalized lymphoblastoid cell line establishment

The study utilized twin family samples from a genetically healthy pedigree, consisting of immortalized lymphoblastoid cell lines (LCLs) derived from the father (CNGB030634, C34), mother (CNGB030632, C32), and their monozygotic (MZ) twin daughters (CNGB030633, C33 and CNGB030635, C35). All participants were of Han Chinese ancestry, representing a genetically homogeneous population characteristic of China. Epstein-Barr virus (EBV) transformation was employed to establish immortalized LCLs from peripheral blood B lymphocytes. Peripheral blood samples and clinical information were obtained after written informed consent was secured from all participants, in compliance with institutional ethical guidelines.

### Genomic DNA extraction

The immortalized lymphoblastoid cell lines were cultured in RPMI 1640 medium (Gibco, 11875119) supplemented with 10% fetal bovine serum (Gibco, A5669701) and maintained at 37°C with 5% CO_2_ in a humidified incubator. The cells were passaged every 3-4 days and harvested after three consecutive passages for cryopreservation.

For cell collection, the cells were washed twice with PBS buffer, followed by DNA extraction using either the BAC-long Blood/Cell DNA Extraction Kit (GrandOmics, XJZZ-BDE-003-1) for ultra-long Oxford Nanopore Technology (UL-ONT) DNA preparation or the Blood and Cell Culture DNA Mini Kit (Qiagen, 13323) for standard DNA extraction. The extracted DNA samples were then subjected to quality control assessments, including evaluation of degradation and contamination by 1% agarose gel electrophoresis, purity measurement using a NanoDrop™ One UV-Vis spectrophotometer (Thermo Fisher Scientific, USA) with acceptable OD260/280 ratios between 1.8-2.0 and OD260/330 ratios between 2.0-2.2, and precise quantification of DNA concentration using a Qubit 4.0 Fluorometer (Invitrogen, USA).

### Sequence data generation

Sequencing data from orthogonal short- and long-read platforms were generated as follows:

### Short reads

Whole-genome sequencing (WGS) datasets were performed on paired-end 150 bp libraries using the MGI-2000RS platform, for the twin daughters (C33 and C35), and their parents (mother C32, and father C34). Raw reads underwent stringent quality control. We excluded reads that met any of the following criteria: length <25 bp; ≥10% unidentified bases (N); ≥50% of bases with Phred quality <5; >10 nt adapter sequence detected (allowing ≤10% mismatches); pronounced sequence-composition bias (A/T to G/C ratio difference >20%); or potential PCR duplicates.

### PacBio HiFi reads

PacBio HiFi data were generated according to the manufacturer’s recommendations. The DNA was used to generate PacBio HiFi libraries using the SMRTbell Prep Kit 3.0. Size selection was performed either with diluted AMPure PB beads according to the protocol, or with Pippin HT using a high-pass cut-off between 10-17 kb based on shear size. Libraries were sequenced on the Revio platform on Revio SMRT Cells and Revio polymerase kit v1 with 2 h pre-extension and 24 h movies on SMRT Link (v.12.0). The library was quantified using a Qubit fluorometer and subsequently sequenced on PacBio Revio systems at GrandOmics, Berry Genomics, and Axbio companies.

### Ultra-long ONT reads

Ultra-high molecular mass gDNA was extracted from the lymphoblastoid cell lines according to a previously published protocol. Libraries were constructed using the Ultra-long DNA Sequencing Kit (ONT, SQK-ULK114) with modifications to the manufacturer’s protocol: ∼40 μg of DNA was mixed with FRA enzyme and FDB buffer as described in the protocol and incubated for 10 min at room temperature, followed by heat inactivation for 10 min at 75°C. RAP enzyme was mixed with the DNA solution and incubated at room temperature for 1 h before the clean-up step. Clean-up was performed using the Nanobind UL Library Prep Kit (Circulomics, NB-900-601-01) and eluted in 450 μL EB. Then, 75 μL of library was loaded onto a primed FLO-PROM114 (R10.4.1) flow cell for sequencing on the PromethION (using MinKNOW software (v.21.02.17-23.04.5), with two nuclease washes and reloads after 24 and 48 h of sequencing. All ONT base calling was done with Dorado (v0.7.1). The sequencing was performed at the GrandOmics Genomics Center in Wuhan, China.

### Reads binning

The long reads were assigned to haplotypes to estimate the depth and assembly accuracy of diploid assemblies. The UL-ONT reads were assigned using Canu (v2.2) haplotype program with parental-specific kmer generated from MGI shotgun reads.

### Generation of phased genome assemblies

Phased genome assemblies were generated using two different algorithms, namely verkko (v2.2.1)^16^ and hifiasm (v0.25.0-r726)^17^ with ONT support. Owing to active development of the verkko and hifiasm algorithms, assemblies were generated with two different versions. Phased assemblies for C33 and C35 were generated using a combination of HiFi and ONT reads using parental short-reads *k-*mers for phasing.

### Contig construction

Initially, the backbone of the diploid genome was constructed using only UL-ONT reads with hifiasm in trio mode, employing the parameters described above. This process generated assemblies of 3.03 Gb for the maternal genome and 3.02 Gb for the paternal genome. Except for chromosomes 13, 14, 15, 21, and 22, the remaining 18 scaffolds were found to constitute complete chromosomes. To further validate the accuracy of the phased contigs, these contigs were assigned to both maternal and paternal genomes because they were homozygous and formed the main path of the graph. Additionally, we utilized other assembly results based on PacBio HiFi and/or ONT reads using verkko and/or hifiasm under trio mode. These results were manually reviewed to confirm the accuracy of the corresponding regions, and any abnormal regions were replaced accordingly.

### Scaffolding

For each haplotype backbone sequence, they were assigned to the respective chromosomes and placed along the coordinates by mapping to the complete version of the human reference genome Telomere-to-telomere (T2T)-CHM13 (v2.0). Of note, CHM13 genome was used to merely help contig placement, but not for reference-guided scaffolding. Within each chromosome, contigs were merged into a longer scaffold based on the overlap between hifiasm and verkko. After that, 22, 18, 22 and 18 gap-free chromosomes were generated in C33 and C35 maternal and paternal phased genomes, respectively, and containing 3, 6, 2 and 6 gaps, respectively. We aligned haplotype sequence to both CN1 v1.0.1 and YAO v2.0 using Minimap2 (v.2.24)^53^. A complete pipeline for this reference alignment is available at GitHub (https://github.com/mrvollger/asm-to-reference-alignment). We also generated a trimmed version of these alignments using the rustybam (v.0.1.33) (https://github.com/mrvollger/rustybam) function trim-paf to trim redundant alignments that mostly appear at highly identical repeats. With this, we aim to reduce the effect of multiple alignments of a single contig over these duplicated regions.

### Gap-filling

Based on the connected chromosomal sequences, TGS-GapCloser (v1.2.1)^18^ was selected as the core gap-closing tool. Other assembly versions of the sequences, along with local assemblies, were employed. Gaps in the sequence were manually filled using binned ONT reads until the gaps were successfully closed. More complex cases were manually examined using IGV (v2.14.1) and JBrowse (v1.16.12), with visualizations created using LINKVIEW (https://github.com/YangJianshun/LINKVIEW).

### Polishing

To minimize introducing switch errors during downstream polishing, the T2T polishing pipeline (https://github.com/arangrhie/T2TPolish/blob/master/doc/T2T_polishing_case_study. md) was adapted to polish the twin genome, primarily using the variants called from binned ONT reads to determine the genotype and guide the selection of the heterozygous variants called from HiFi and short reads. Long reads (binned ONT and all HiFi data) were mapped using Winnowmap2, and primary alignments were retained. MGISEQ short reads were mapped using BWA-MEM2 (v2.3)^54^, and duplications were marked using “bamsortmarkdup” from biobambam2 (v2.0.183). Small variants were called using the “hybrid” model in GATK (v4.3.0.0)^55^ based on the combined HiFi and MGISEQ alignments. Variants marked as ‘PASS’ by deepvariant with VAF ≥ 0.5 were retained, and the called genotype was required to be ‘1/1’. These variants were further filtered using merfin and used to polish the maternal and paternal genomes.

Based on the assembly-only kmer coordinates reported by merqury (v1.3)^20^, the binned HiFi and ONT read alignment at the region was further visualized using the modified bamsnap for manual examination. The variants called in the regions were further manually examined and used to polish the genome with nextpolish2 (v0.2.1).

Structural variants were called based on the binned ONT alignment using Sniffles2 (v2.6.3)^56^. The region from both binned ONT alignment and unbinned HiFi alignment we extracted based on the ONT alignment and visualized using a modified version of bamsnap (https://github.com/zy041225/bamsnap). The ONT and HiFi alignment was further manually examined to confirm the precise coordinate and size of each structural variation (SV). These small variants and large SVs were then merged and used to polish the genome using merfin (v1.1)^57^ and bcftools (v1.16).

These variants and polish rounds were combined following the workflow shown in Supplementary Figure.

### Evaluation of phased genome assemblies

To evaluate the base pair and structural accuracy of each phased assembly, we used a multitude of assembly evaluation tools as well as orthogonal datasets such as PacBio HiFi, ONT and short-read data. We note that we fixed four haplotype switch errors in our assembly-based variant callsets to avoid biases in subsequent analysis.

### Read to assembly alignment

To evaluate *de novo* assembly accuracy, we aligned sample-specific PacBio HiFi reads to their corresponding phased genome assemblies. HiFi and ONT read sets were aligned to the diploid genome (all-to-dip) as well as to each haploid genome (all-to-hap) with Winnowmap (v2.03) or Minimap2 (v2.29-r1283) using the pipeline from T2T-Polish (https://github.com/arangrhie/T2T-Polish/tree/master/winnowmap).

### Genome Continuity Inspector (GCI) validation

GCI (v1.0)^21^ is an assembly assessment tool for high-quality genomes (e.g. T2T genomes), in base resolution. After stringently filtering the alignments generated by mapping long reads (PacBio HiFi and/or ONT long reads) back to the genome assembly aligned using Winnowmap (v.2.03) and Minimap2 (v2.29-r1283). GCI will report potential assembly issues and also a score to quantify the continuity of assembly. GCI was used to identify regions of collapse. The regions where the depth exceeded the Minimum and Maximum depth in folds of mean coverage, as well as areas of misassembly where all nucleotide counts were zero, were identified.

The binned HiFi and binned ONT reads were mapped to the corresponding maternal (or paternal) genome. Primary alignments from each aligner were separately extracted and processed following the method described at https://github.com/arangrhie/T2T-Polish/tree/master/coverage. The regions were further merged based on the three HiFi (or ONT) results, and only the region was retained if one or more alignments obtained using Minimap2 and Winnowmap2 suggested a potential problem. The read alignment was further confirmed using the modified bamsnap and IGV to exclude false positives introduced by read binning. The problematic HiFi and ONT regions were further combined only when one region showed abnormal coverage in both HiFi and ONT alignment. Most potentially problematic regions were in the centromeres. Only the regions outside the centromeric regions were examined and confirmed.

### Assembly base-pair quality value (QV)

To evaluate the accuracy of the genome assembly, PacBio HiFi reads and short pair-end reads were utilized to construct a hybrid kmer dataset and evaluate the assemblies using Meryl (v.1.4.1) to count the *k-*mers of length 21 and 31. We then used Merqury (v.1.3), which compares the *k-*mers from the sequencing reads against those in the assembled genome and flags discrepancies where *k-*mers are uniquely found only in the assembly. These unique *k-*mers indicate potential base-pair errors. Merqury then calculates the QV based on the *k-*mer survival rate, estimated from Meryl’s *k-*mer counts, providing a quantitative measure to assess the completeness and correctness of the genome assembly.

### Gene annotation

Gene annotation was performed by employing liftoff (v1.6.3)^58^ and LiftOver (v438)^59^ to project the T2T-CHM13v2.0 reference annotation onto the assembly as the main source. Moreover, human uniprot_sprot (release-2022_05) protein data using miniprot (v0.17)^60^, BRAKER (v3)^61^ annotation, and Augustus (v3.4.0)^62^ annotation were integrated into the gene dataset using maker3 (v3)^63^. The obtained annotation was further complemented with tRNA annotation using tRNAscan-SE (v2.0)^64^ and other ncRNA annotation using Rfam (v14.9)^65^.

### Analysis of subtelomeric sequence composition

We identified segmental duplications (SD) and subtelomeric repeats (Srpt) within the terminal 500 kb of each chromosome in both C33 and C35 haplotype assemblies. Srpt was defined as ≥1 kb blocks showing ≥90% identity to other subtelomeric sequences, whereas SD^66^ was assigned to ≥1 kb blocks with ≥90% identity to any region of the whole-genome, and regions that met both rules were simply called SD-Srpt. RepeatMasker run with high-sensitivity parameters was used to annotate interspersed repeats and interstitial (TTAGGG)n-like tracts. Each assembly was oriented from telomere to centromere, and self-BLAST (v2.17)^67^ searches were performed to exclude artificial duplications. The output generated by this integrated pipeline was used to quantify the total amount of segmental duplications and subtelomeric repeats. The extent of enrichment compared to the genome-wide average was also assessed.

### Satellite and repeat annotations

To identify canonical and novel repeats, we utilized the previously described pipeline, with modifications to include both the Dfam (v3.691)^68^ and Repbase (v20181026) libraries for each species during RepeatMasker annotation. An initial RepeatMasker run identified canonical repeats, while a subsequent RepeatMasker run was completed to include repeat models first identified in the analysis of T2T-CHM13^5^ and T2T-Y^69^. To identify and curate previously undefined satellites, we utilized additional TRF^70^ and ULTRA screening of annotation gaps >10 kbp in length. Potential gaps were identified via BEDTools (v2.29.083)^71^ by subtracting both the repeat and gene annotations for sequence. To identify potential redundancy, satellite consensus sequences generated from gaps identified in each genome were compared using crossmatch and were used as a RepeatMasker library to search for overlap in the other human T2T genome. Consensus sequences were considered redundant if there was a significant annotation overlap in the RepeatMasker output. Repeat consensus sequences were manually curated using RepeatMasker searches to ensure accuracy and identify additional variants.

### Genomic comparison

After masking the centromeric and heterochromatic regions into Ns, each chromosome of C33/C35 maternal (or paternal) genome was aligned to the corresponding chromosome of CHM13 using nucmer (v4.0.0rc1)^72^ with parameter set “--maxmatch -t 12 -l 100 -c 500”. Delta results were later filtered using delta-filter with parameter “-m -i 90 -l 100”, and SVs were identified using SyRI (v1.6.3)^73^. Only SVs >50 bp were retained. These SVs, along with the flanking regions, were then manually examined using LINKVIEW to filter out false-positive SVs introduced due to problematic alignment. The SVs of the combined C33/C35 genome were generated by combining the SVs detected between CHM13 and C33/C35.Mat genome and between CHM13 and C33/C35.Pat genomes.

### 5mC methylation, differentially methylated cytosine (DMC) and differentially methylated region (DMR) detection

We base-called and aligned ONT reads with Dorado and Minimap2, then applied DeepMod2^74^ to extract 5mC methylation probabilities at each cytosine guanine dinucleotide (CpG). Per-read scores were averaged across replicates to build a single matrix of methylation frequencies for maternal and paternal haplotype of C33 and C35. This matrix was supplied to metilene^75^ for differentially methylated region calling with tmin = 10 and a minimum mean difference of 0.1. Significant DMRs were identified using the adjusted two-dimensional Kolmogorov-Smirnov *p*≤0.05, yielding a comprehensive set of regions capable of discriminating between twins. The differential CpG methylation analysis using Fisher’s exact test on proportion value with CpGtools^76^.

Additionally, imprinting is defined as the preferential expression of one parental allele over the other, primarily regulated by differential methylation at CpG dinucleotides. Here, we utilized a set of validated imprinting-associated DMRs defined by Zink et al^77^, and assessed the concordance of methylation patterns in our twin pairs across these regions with parent-of-origin (PofO)-DMR scores > 25.

### Centromere identification and annotation

To identify the centromeric regions within each haplotype genome, we first aligned the whole-genome assemblies to the T2T-CHM13 v2.0 reference genome using Minimap2 (v2.24) with the following parameters: -I 10G -a --eqx -x asm20 -s 5000. We filtered the alignments to only those regions that traversed each human centromere, from the p- to the q-arm, using SAMtools (v1.9) and then ran RepeatMasker (v4.1.0) to identify regions containing α-satellite sequences, marked by “ALR/Alpha”. Once we identified the regions of the assemblies containing α-satellite repeats, we ran Snakemake-HumAS-SD (https://github.com/logsdon-lab/Snakemake-HumAS-SD/tree/main) with AS-higher-order repeats (HORs)-hmmer3.4-071024.hmm Hidden Markov Model. This generated a BED file with each α-satellite suprachromosomal family (SF) designation and its organization along the centromere. We used the α-satellite SF BED file to visualize the organization of the α-satellite HOR arrays with R (v1.1.383) and the ggplot2 package. Moreover, Inter-chromosomal centromere alignments were executed with RaMA^78^, an aligner optimized for ultra-long tandem repeats, ensuring base-level accuracy across highly divergent satellite arrays.

We validated the construction of each centromeric region by first aligning native PacBio HiFi and ONT data from the same source genome to each whole-genome assembly using Minimap2 or Winnowmap (v2.03)^79^ (for ONT data). We, then, assessed the assemblies for uniform read depth across the centromeric regions via IGV and for collapses, duplications, and misjoins via NucFreq^80^. Centromeres that were found to have a misassembly were flagged and are indicated in the figures.

To generate pairwise sequence identity heatmaps of each centromeric region, we ran StainedGlass (v6.7.0)^29^ with the following parameters: window = 5,000, mm_f = 30,000, and mm_s = 1,000.

### Detection of small *de novo* variants

Following the parameters outlined previously, we called variants in HiFi and short read data aligned to C33.Mat haplotype assembly using Clair3 (v1.2.0)^81^ and GATK HaplotypeCaller (v4.3.0.0)^82^. We separated single-nucleotide variant (SNV) and indel calls and applied basic quality filters, such as removing clusters of three or more SNVs in a 1 kb window. We combined this set of variant calls generated by a secondary calling method (https://github.com/Platinum-PedigreeConsortium/Platinum-Pedigree-Inheritance/blo b/main/analyses/Denovo.md) and subjected all calls to the following validation process. We validated both SNVs and indels by examining them in HiFi, ONT and short read data, excluding reads that failed to reach the mapping quality (59 for long reads, 0 for short reads) thresholds. Reads with high base quality (>20) and low base quality (<20) at the variant site were counted separately. We retained variants that were present in at least two types of sequencing data for the child, and absent from high-quality parental reads. For SNV calls, we next examined HiFi data for every sample in the pedigree. We determined an SNV to be truly *de novo* if it was absent from a family member that wasnot a direct descendant of the *de novo* sample. Finally, we examined the allele balance of every variant, determined which variants were in TRs, and re-evaluated parental read data across all sequencing platforms, removing variants with noisy sequencing data or more than two low-quality parental reads supporting the alternative allele.

Considering each sequencing technology individually, we determined that a variant was truly *de novo* if it was present in a child and absent from its parents, and inherited if a parent had reads with the alternate allele.

## Ethical statement

The study was conducted in compliance with ethical standards and received formal approval. All methods were performed in accordance with relevant guidelines and regulations. Written informed consents were obtained from the participants.

## Data Availability

The raw sequencing data of the Chinese family have been deposited in the Genome Sequence Archive for human^83^ at the NGDC, BIG, CAS / CNCB (GSA-Human: HRA013168), and are publicly accessible at https://ngdc.cncb.ac.cn/gsa-human. The C33 and C35 genome sequences have been deposited in the Genome Warehouse^84^ at the NGDC, BIG, CAS / CNCB (GWH: GWHGQMV00000000.1, GWHGQMW00000000.1, GWHGQMX00000000.1 and GWHGQMY00000000.1), and are publicly accessible at https://ngdc.cncb.ac.cn/gwh.

## Acknowledgments

This work was supported by grants from the National Key Research and Development Program of China (2024YFC3405701 to Y.T., 2025YFC3410300 to C.Y. and D.W., 2022YFC3400304 to J.H.) and the National High Level Hospital Clinical Research Funding (BJ-2025-177 to G.L. and K.Z.). We gratefully acknowledge the sequencing services provided by GrandOmics, Berry Genomics, Axbio, MGI and Qitan Technology for this study.

## Author contributions

J.H., C.Y. and K.Z. conceived and designed the study. T.L., L.Q., X.J. and L.C. performed sample preparation and sequencing. D.F. and C.Y. performed the genome assembling. D.F., T.L., G.L., Z.J., Y.Q. and Y.T. curated the data and data analysis. D.W. provided analytical suggestions. T.L. drafted the original manuscript. J.H., C.Y. and D.F. reviewed and edited the manuscript. J.H. and C.Y. supervised the project. All authors read and approved the final version of the manuscript.

## Competing interests

Dongming Fang, Lingxin Qiu, Xin Jin, Lei Cheng and Chentao Yang are employees of MGI Tech. All authors declare no competing interests.

## Notes

https://ngdc.cncb.ac.cn/gwh

### Competing Interest Statement

The authors have declared no competing interest.

