## Supplemental Figures for "Complete and haplotype-resolved maps of genomic and epigenetic discordance in monozygotic twins"

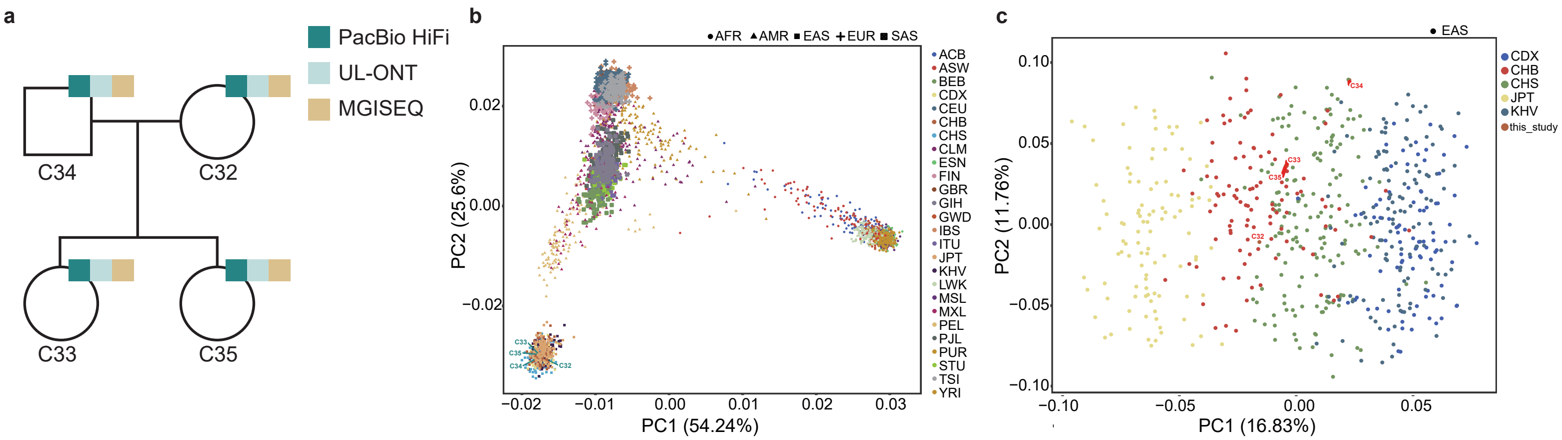


**Supplementary Fig. 1 Schematic illustration of sequencing platforms and population genetic analysis of the MZ family. a** Schematic diagram summarized the three sequencing technologies (PacBio HiFi, UL-ONT, and MGISEQ) applied to the family samples. **b** PCA illustrated the genetic relationship between the four family members and the 1KG populations. **c** PCA focused on Asian populations, where the four family samples are highlighted in a distinct color.

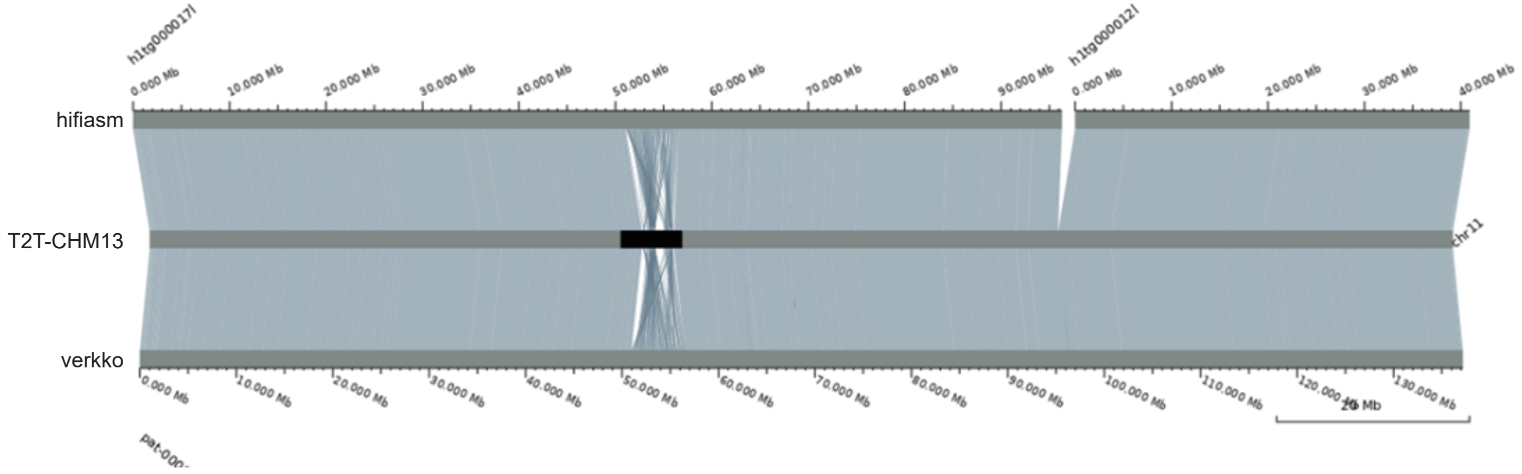


**Supplementary Fig. 2 Illustrations of the merging hifiasm and verkko assemblies.** For the paternal C33 chromosome 11, hifiasm and verkko contigs were aligned to CHM13 (middle panel) to determine their order and orientation. The black block on CHM13 indicated the centromeric region.

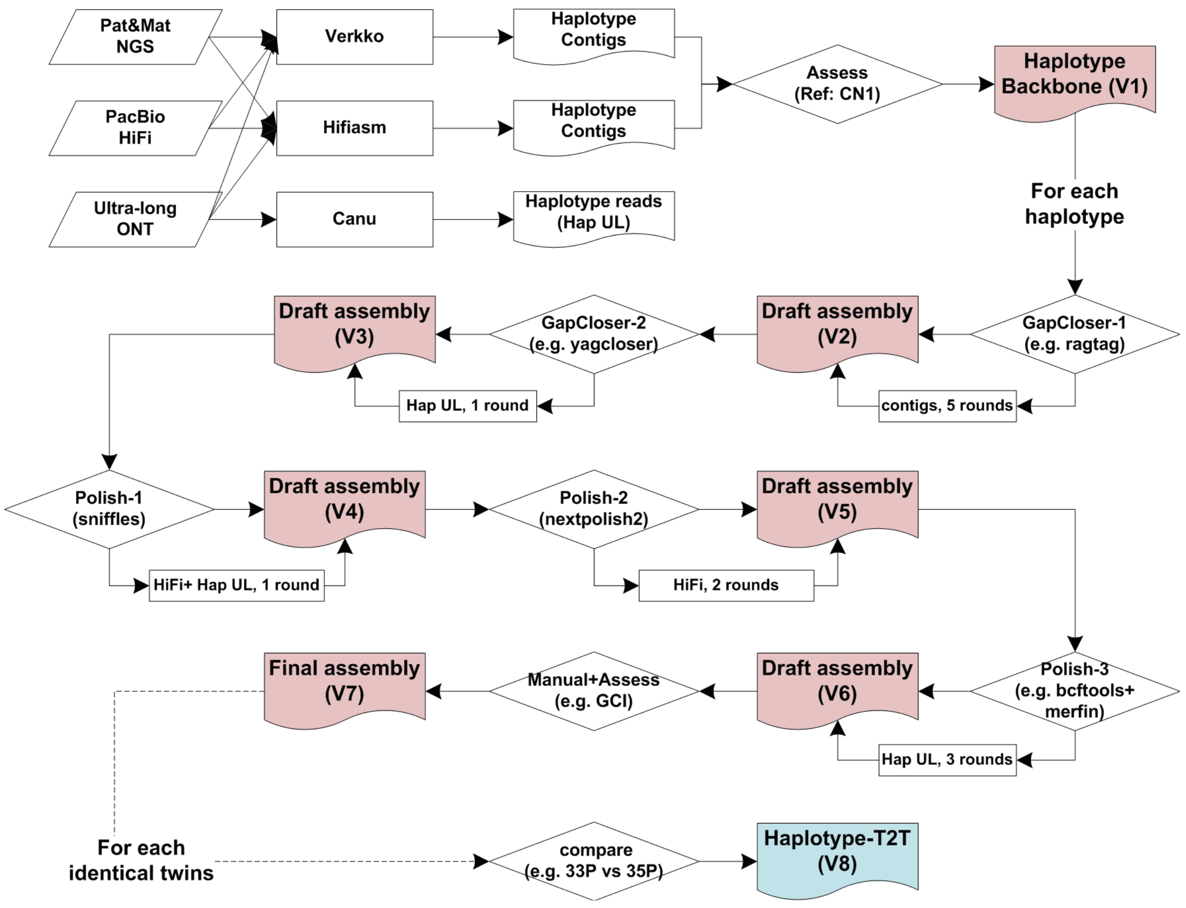


**Supplementary Fig. 3 Genome-assembly workflow for the monozygotic twins C33 and C35.** We devised a haplotype-resolved, T2T assembly pipeline tailored for the monozygotic twins C33 and C35. Short reads from both parents were used to label contigs from hifiasm and verkko assembly with HiFi and UL-ONT, yielding preliminary paternal and maternal bins. With the same parental labels, Canu was employed to partition ONT reads into the corresponding haplotypes, creating phased "ultra-long" read pools for gap closure and polishing. The contigs and Hap UL reads were aligned to the CN1 reference. Coverage and phase consistency were evaluated to select the best haploid backbone for each chromosome. Two-step gap filling: GapCloser-1 (V2): five iterative rounds with RagTag; GapCloser-2 (V3): one refinement round with YagCloser. Three-round polishing: Polish-1 (V4): Sniffles to correct large SVs; Polish-2 (V5): NextPolish2 using HiFi depth for base accuracy; Polish-3 (V6): bcftools + Merfin coupled with Hap UL reads for final base-quality tuning. At every round, we run GCI and Merqury to assess coverage and error region around the whole genome, cumulatively defining the final V7 release. After gap-closing and polishing all four haplotypes (C33.Mat, C35Mat, C33.Pat, C35.Pat), we performed pairwise comparisons (e.g., C33.Pat vs C35.Pat) to purge residual discrepancies, yielding two high-quality, twin-specific T2T haploid genomes.

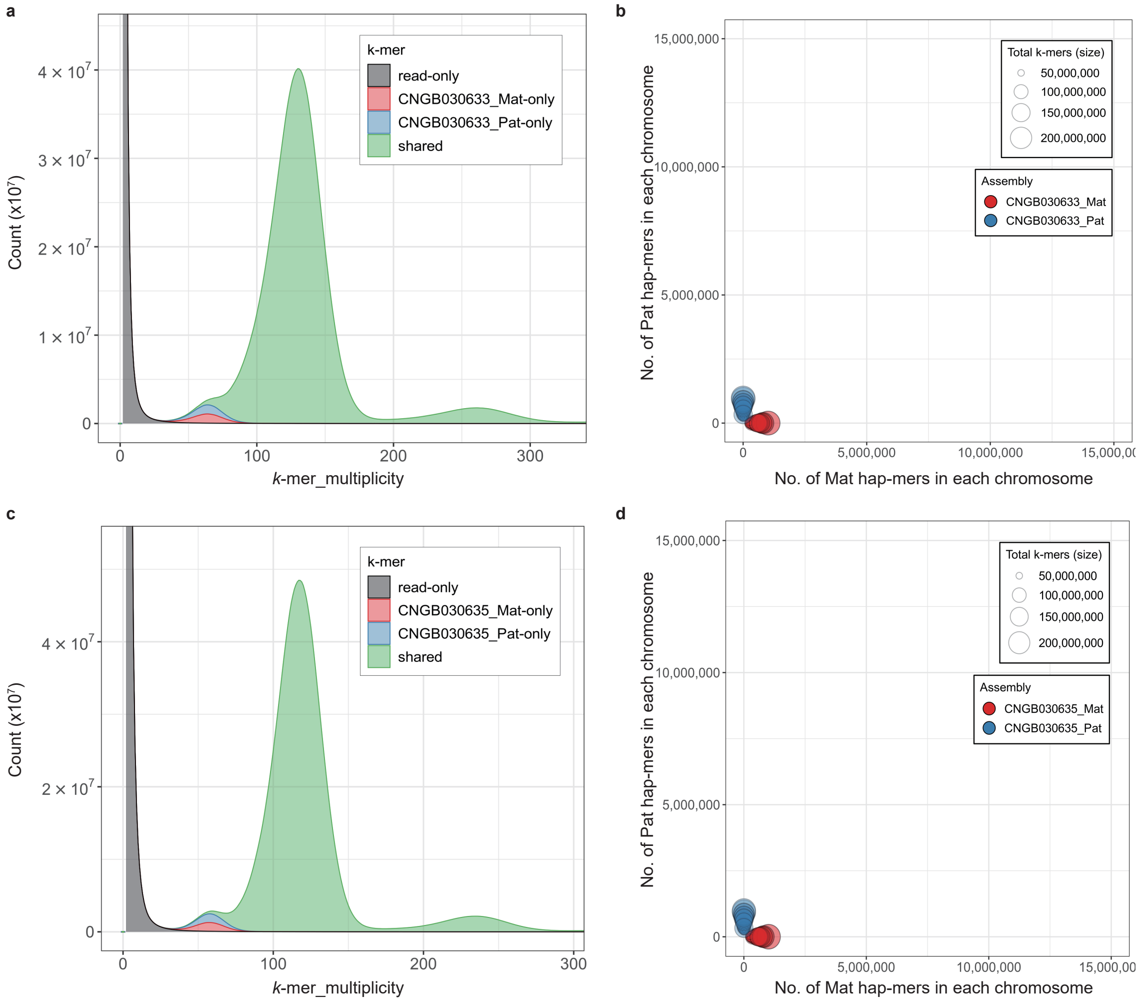


**Supplementary Fig. 4 The quality of the diploid C33 and C35 assembly. a** Spectrum plots of *k*-mers in the assembly and reads used to assess C33 assembly error and completeness. The distributions of *k*-mers found in the reads and in Mat/Pat assemblies were shown in distinct colors. The X-axis represented the *k*-mer_multiplicity, which meant the number of times a *k*-mer occurs in the read set. The *k*-mers at the left end of the X-axis with zero read multiplicity represented erroneous *k*-mers present only in assemblies. Read-only indicated *k*-mers found only in reads. Shared indicated *k*-mers shared in both C33.Mat and C33.Pat assemblies. **b** Hap-mer blob plot of the C33 assembly. Mat and Pat assemblies were colored red and blue, respectively. For each chromosome, Mat and Pat hap-mers were shown as colored blobs whose area was proportional to the chromosome's total *k*-mer count. **c** Spectrum plots of *k*-mers in the assembly and reads used to assess C35 assembly error and completeness. The distributions of *k*-mers found in the reads and in Mat/Pat assemblies were shown in distinct colors. The X-axis represented the *k*-mer_multiplicity, which meant the number of times a *k*-mer occurs in the read set. The *k*-mers at the left end of the X-axis with zero read multiplicity represented erroneous *k*-mers present only in assemblies. Read-only indicates *k*-mers found only in reads. Shared indicates *k*-mers shared in both C35.Mat and C35.Pat assemblies. **d** Hap-mer blob plot of the C35 assembly. Mat and Pat assemblies were colored red and blue, respectively. For each chromosome, Mat and Pat hap-mers were shown as colored blobs whose area was proportional to the chromosome's total *k*-mer count.

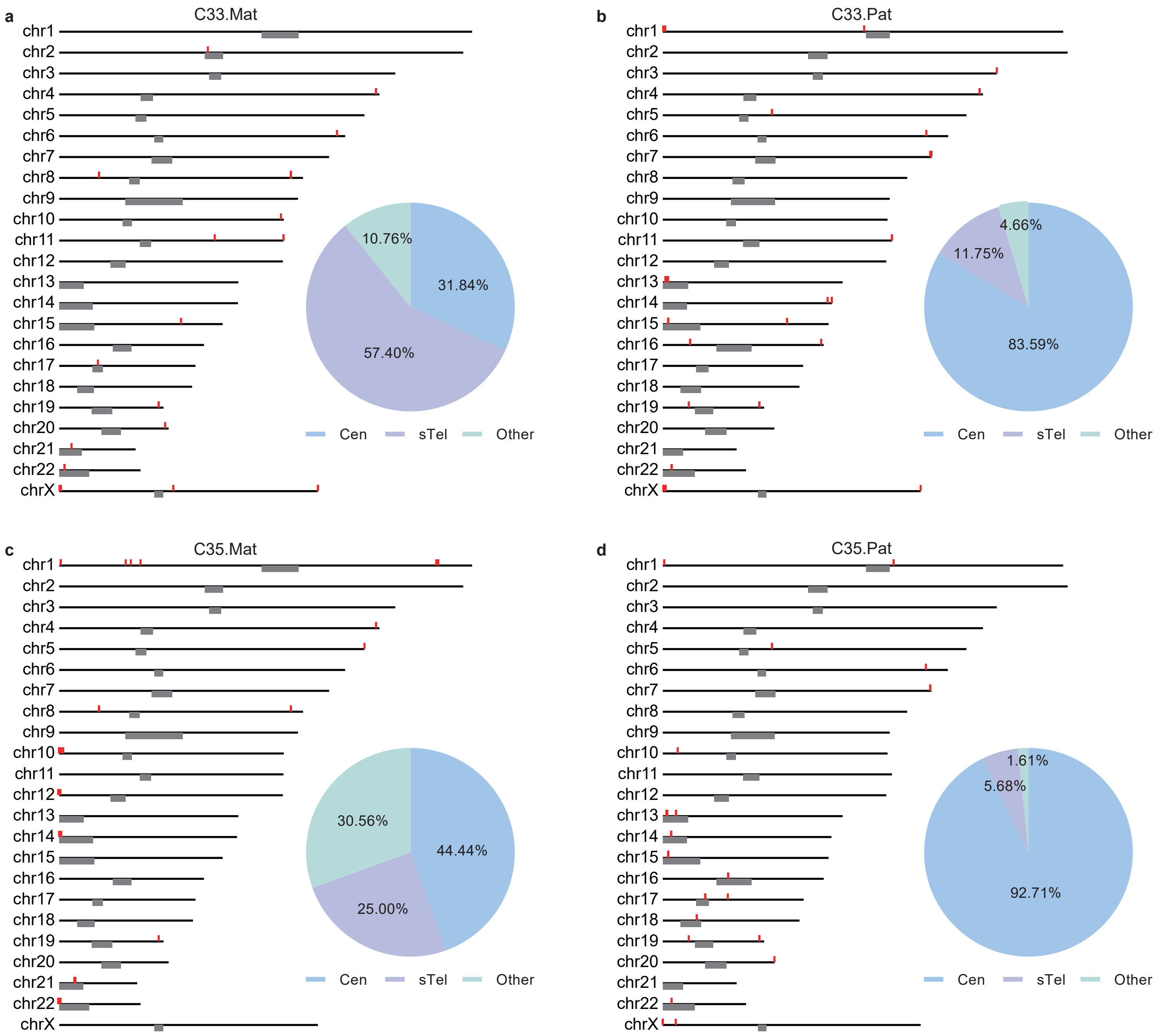


**Supplementary Fig. 5 Evaluation of erroneous *k*-mers.** Evaluation of erroneous *k*-mers of C33.Mat (a), C33.Pat (b), C35.Mat (c) and C35.Pat (d), and gray bars mark the centromeres of each chromosome. Cen, centromere. sTel, subtelomeres.

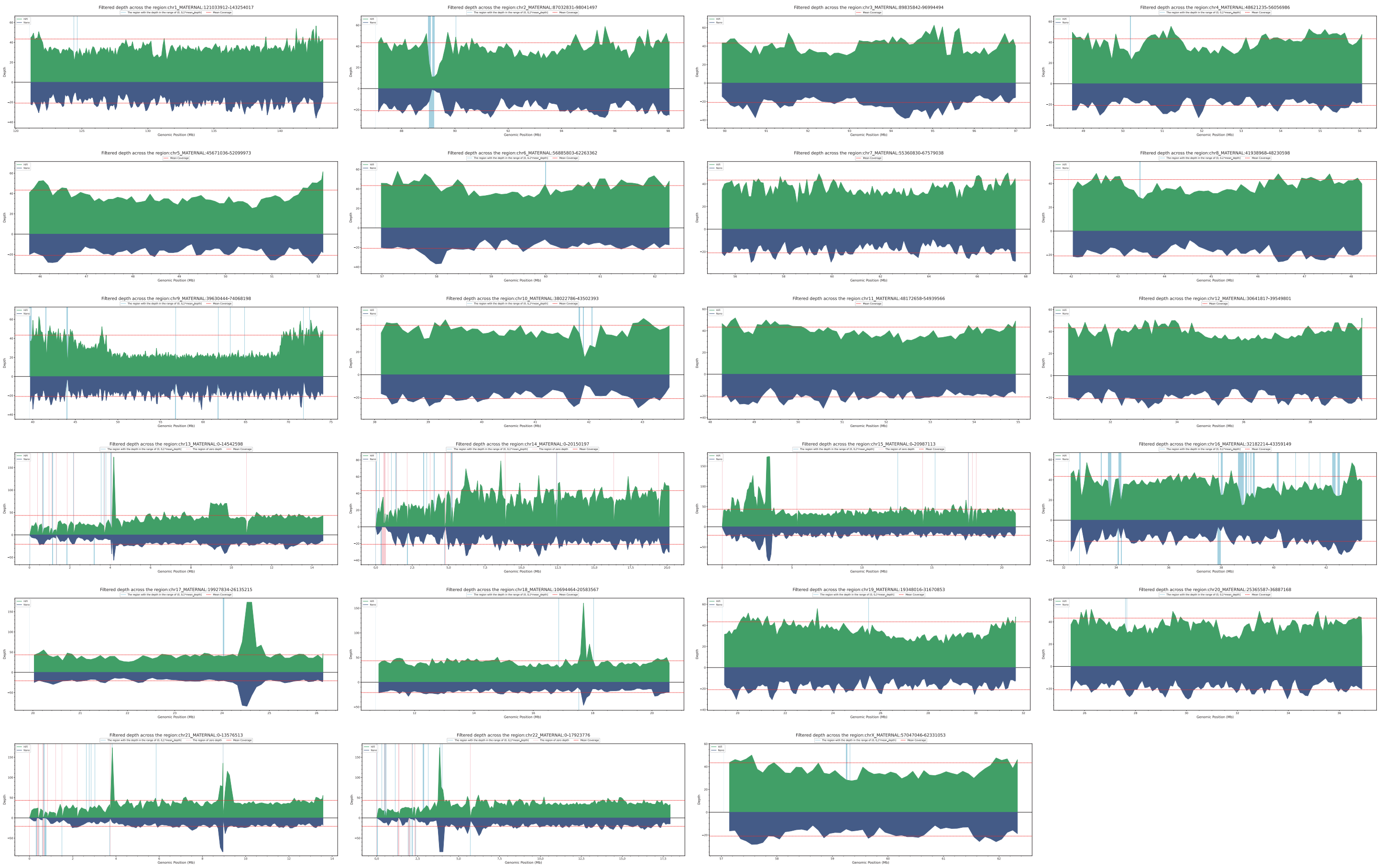

**Supplementary Fig. 6 GCI plot of binned ONT and HiFi reads of C33.Mat centromere region.** GCI, genome continuity inspector.

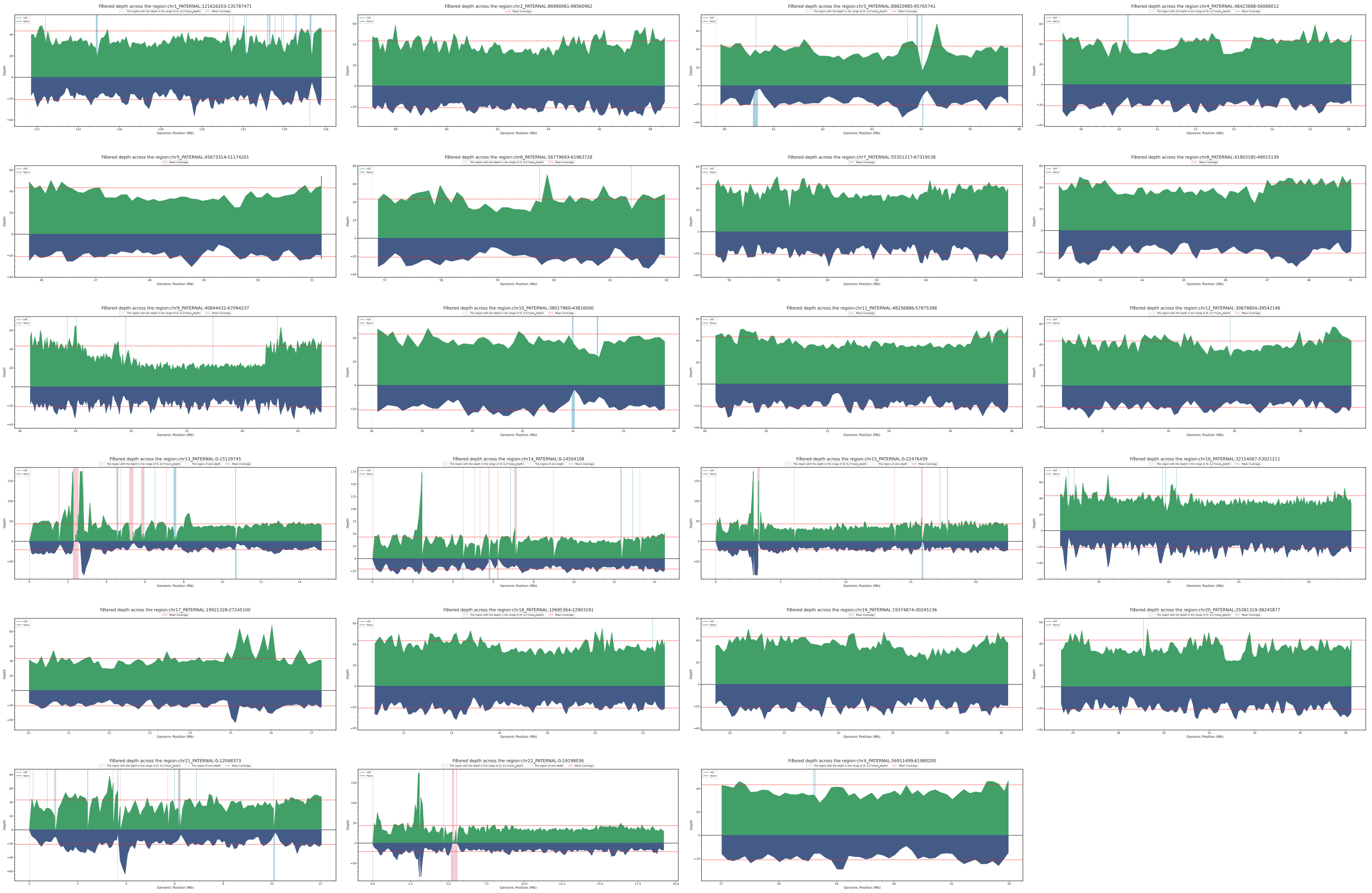

**Supplementary Fig. 7 GCI plot of binned ONT and HiFi reads of C33.Pat centromere region.** GCI, genome continuity inspector.

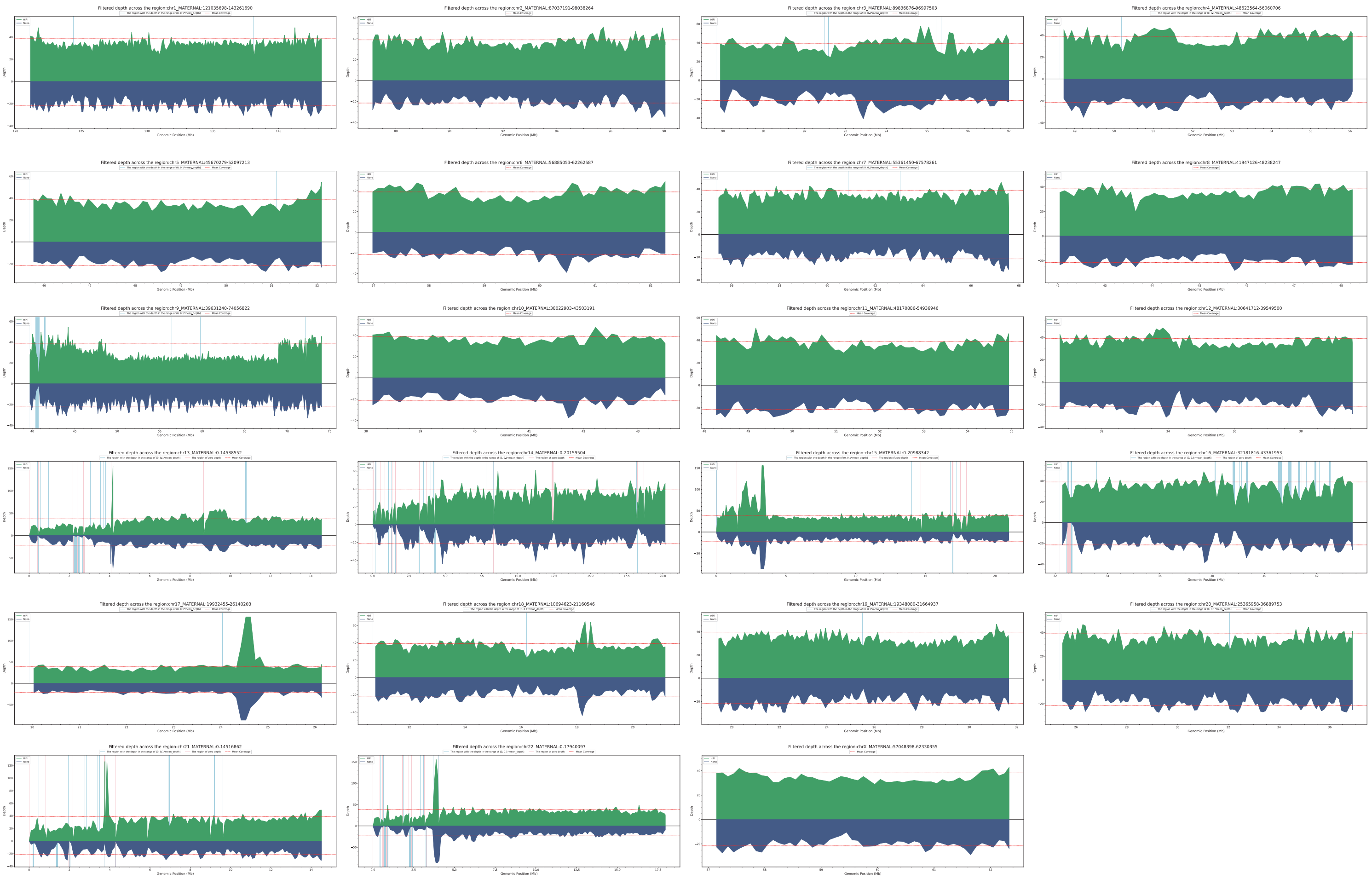

**Supplementary Fig. 8 GCI plot of binned ONT and HiFi reads of C35.Mat centromere region.** GCI, genome continuity inspector.

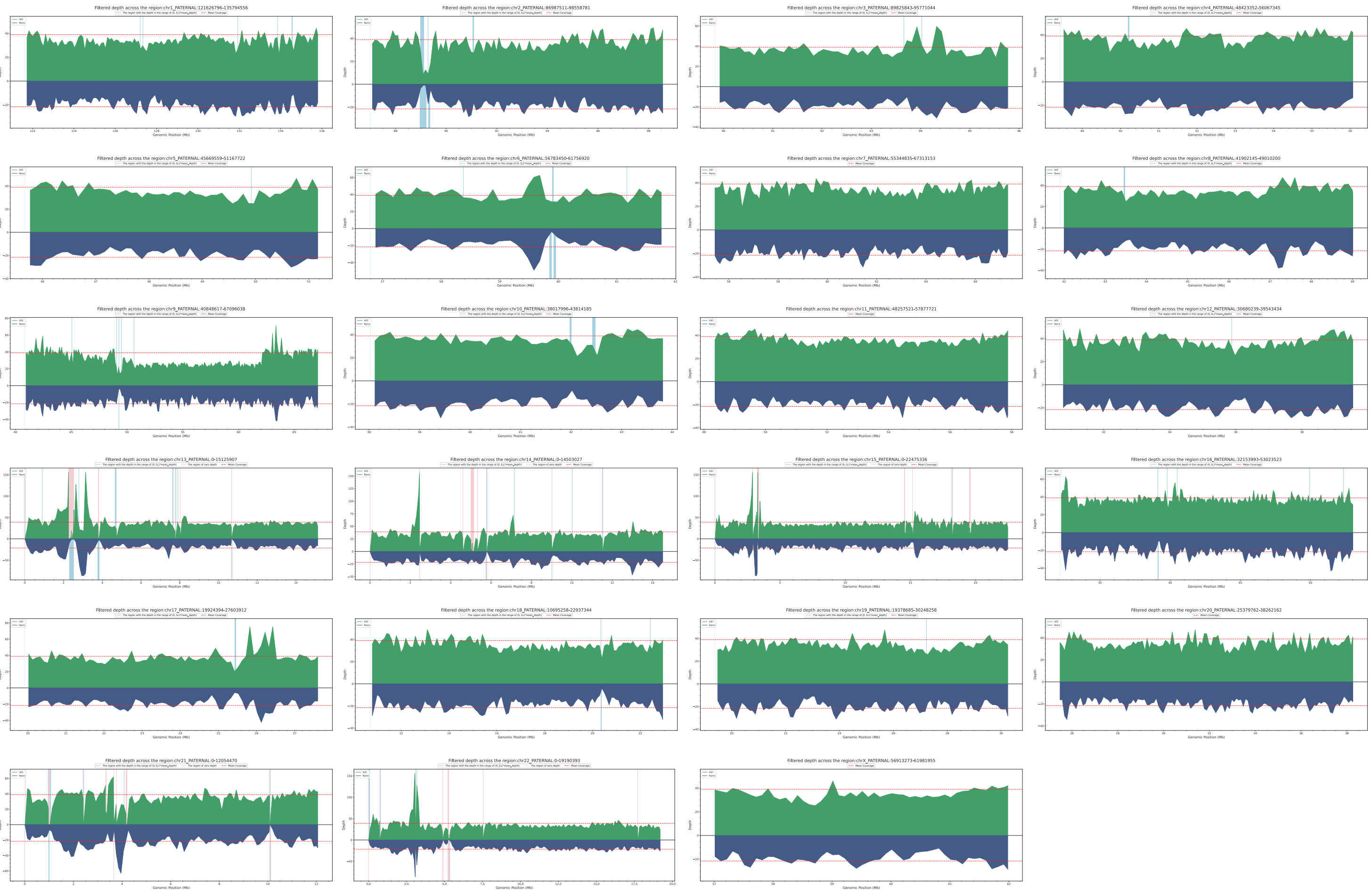

**Supplementary Fig. 9 GCI plot of binned ONT and HiFi reads of C35.Pat centromere region.** GCI, genome continuity inspector.

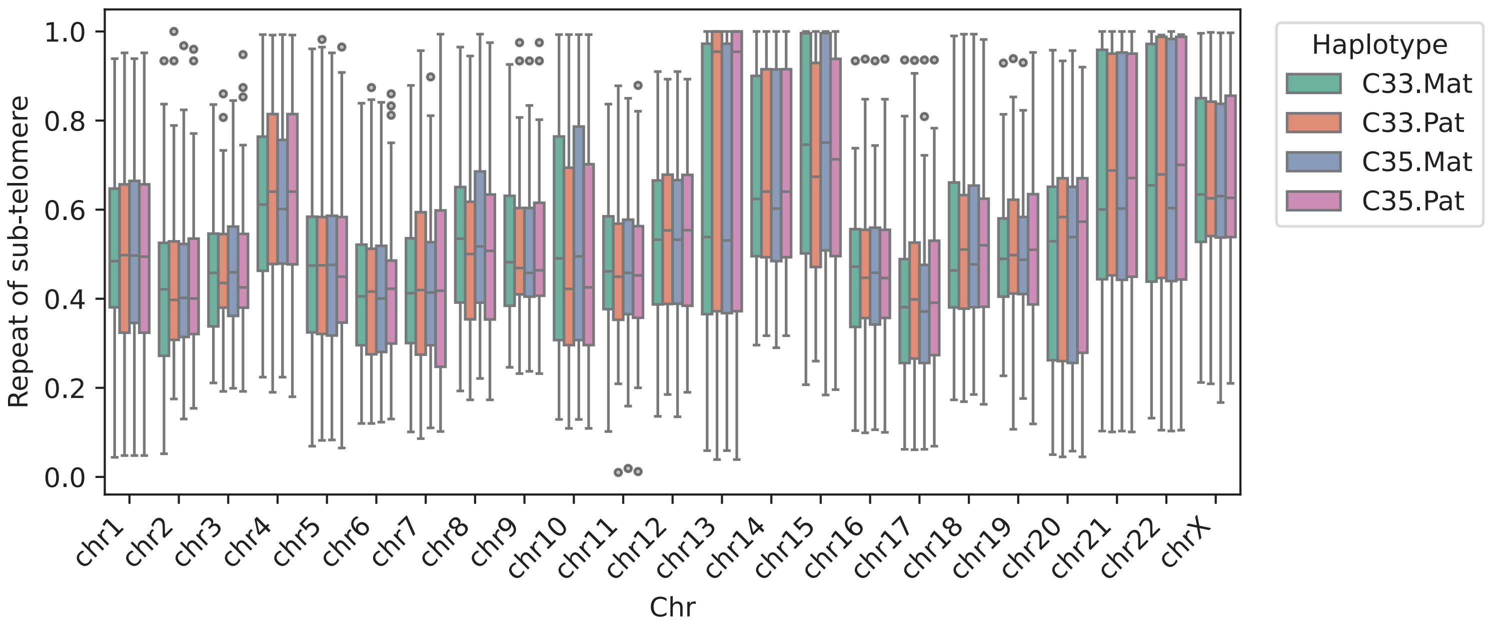


**Supplementary Fig. 10 Subtelomeric repeat landscape of the four haplotypes (C33.Mat, C33.Pat, C35.Mat, C35.Pat).** For each chromosome, the terminal 500 kb was divided into 10 kb windows and the proportion of bases annotated as repetitive elements was analysed. Repeat density rised sharply toward the telomere, with acrocentric short arms (e.g., chromosome 13, 14, 15, 21, 22) reaching >80%, while chromosomes 2, 6, 7 and 17 remained below 40%, reflecting chromosome-specific expansion and contraction of terminal repeat arrays.

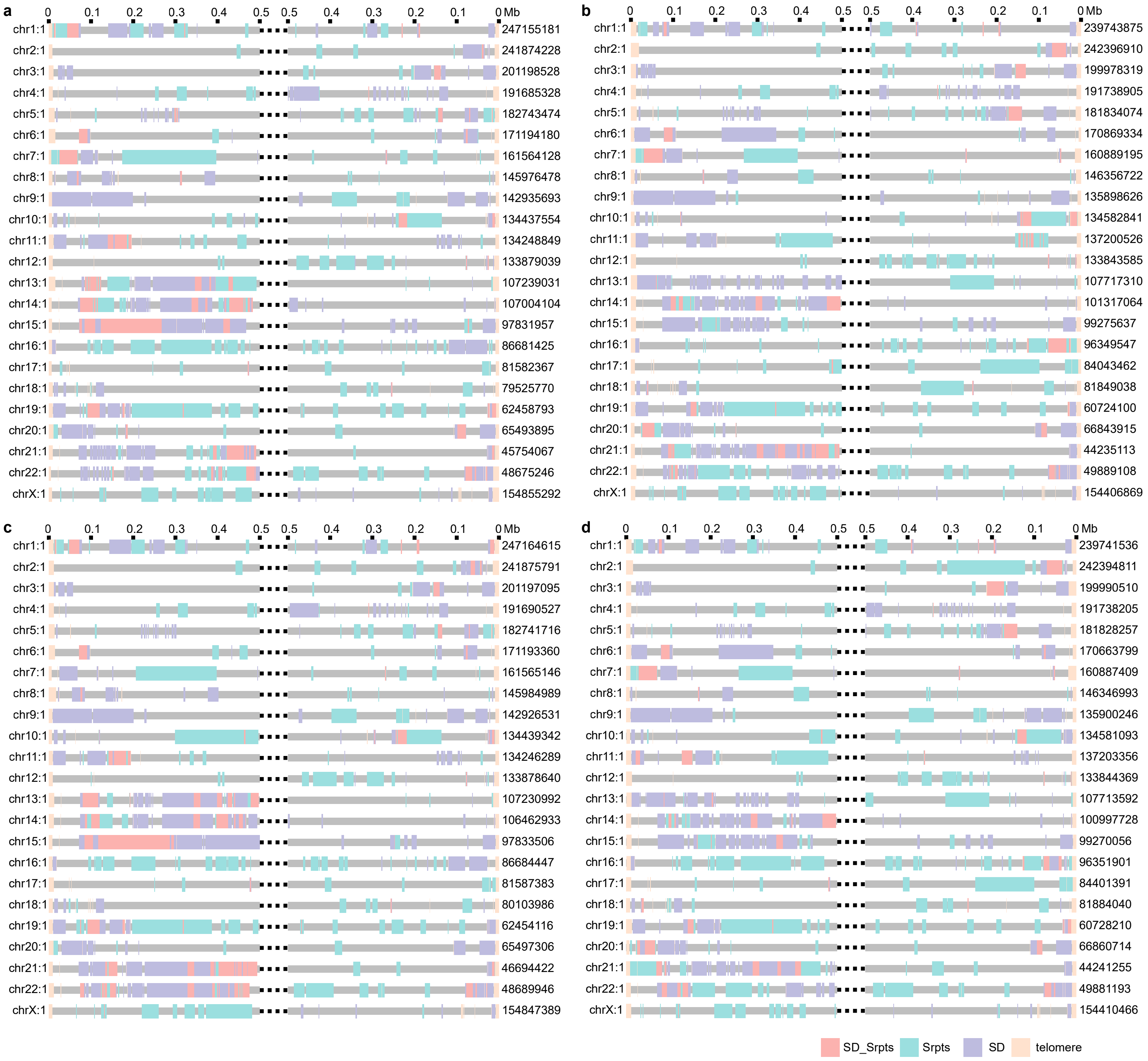


**Supplementary Fig. 11 Duplications in each chromosome terminal 500 kb of the four haplotypes (C33.Mat (a), C33.Pat (b), C35.Mat (c), C35.Pat (d)).** SD and Srpts were both >1 kb duplication events. Their cumulative lengths were plotted as stacked bars.

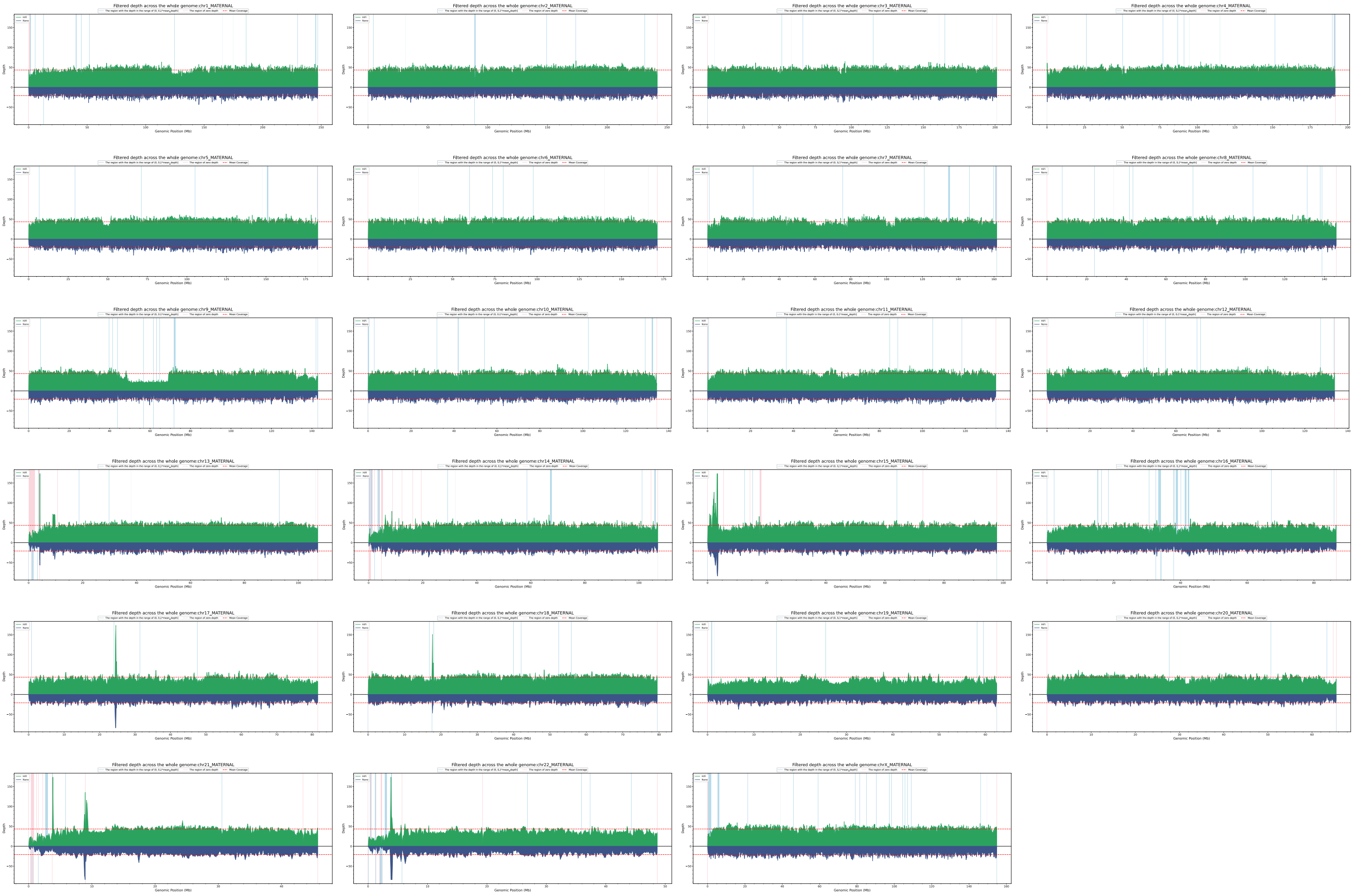


**Supplementary Fig. 12 GCI plot of binned ONT and HiFi reads along the C33.Mat's genome.** GCI, genome continuity inspector.

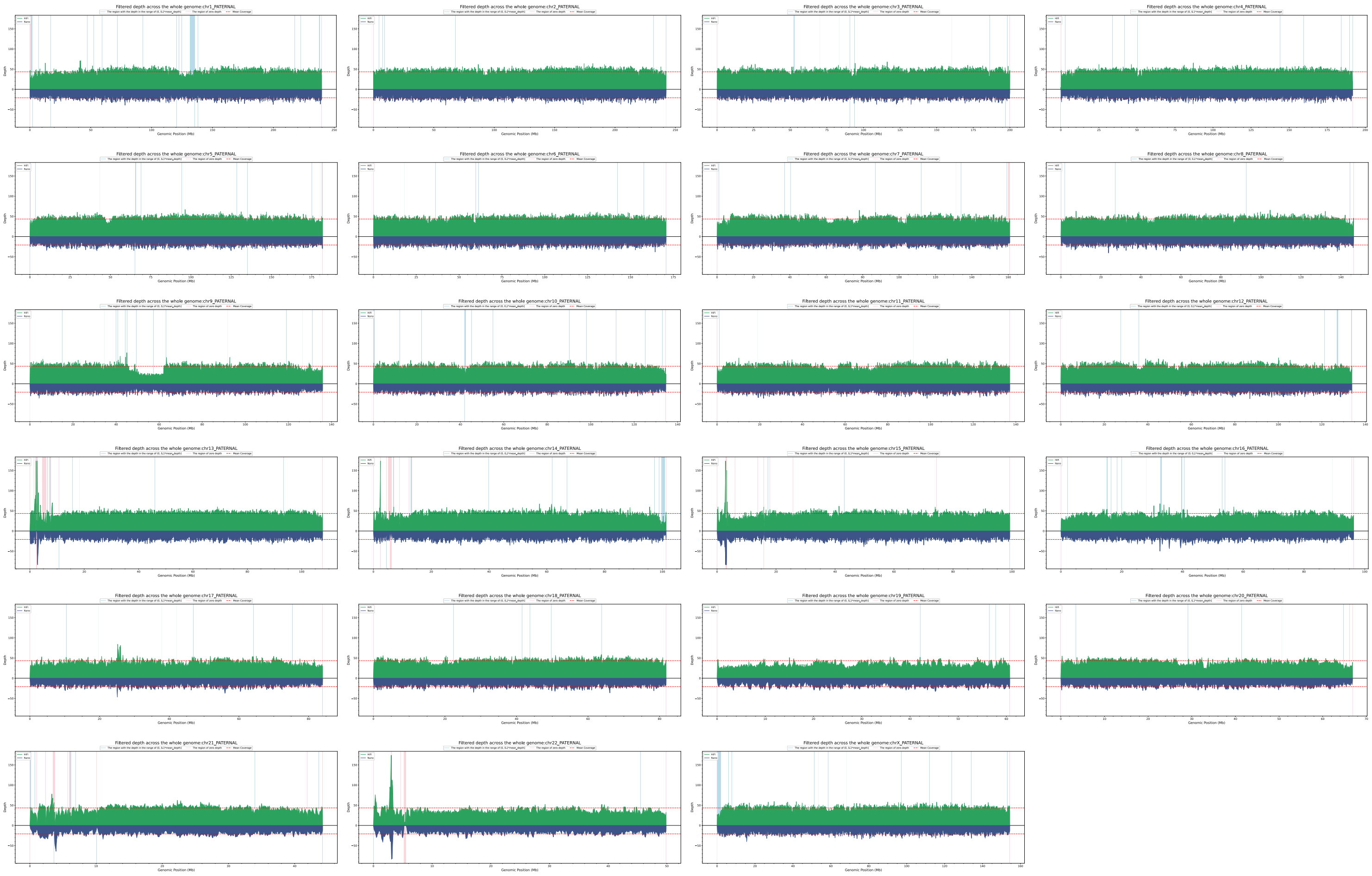

**Supplementary Fig. 13 GCI plot of binned ONT and HiFi reads along the C33.Pat's genome.** GCI, genome continuity inspector.

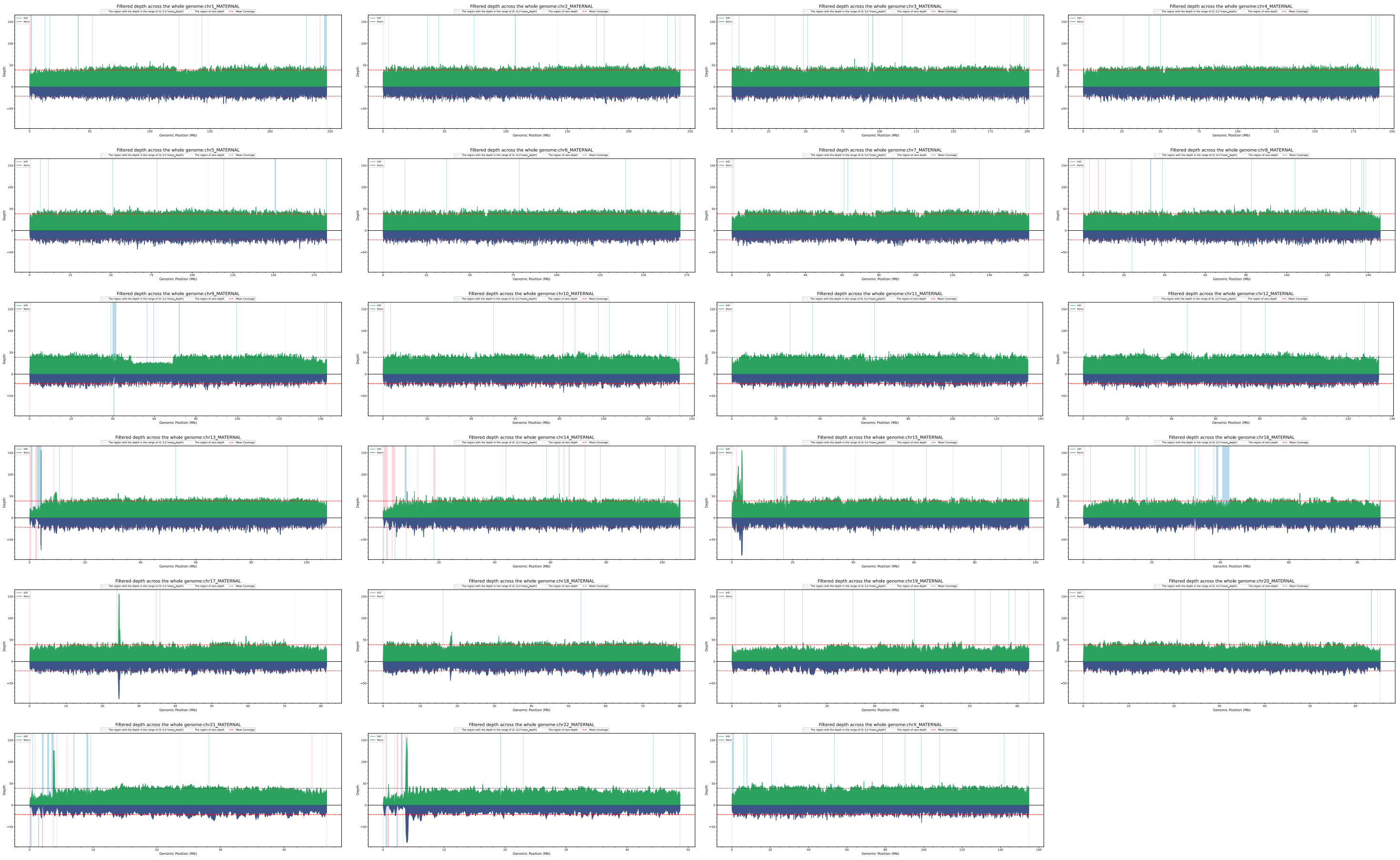


**Supplementary Fig. 14 GCI plot of binned ONT and HiFi reads along the C35.Mat's genome.** GCI, genome continuity inspector.

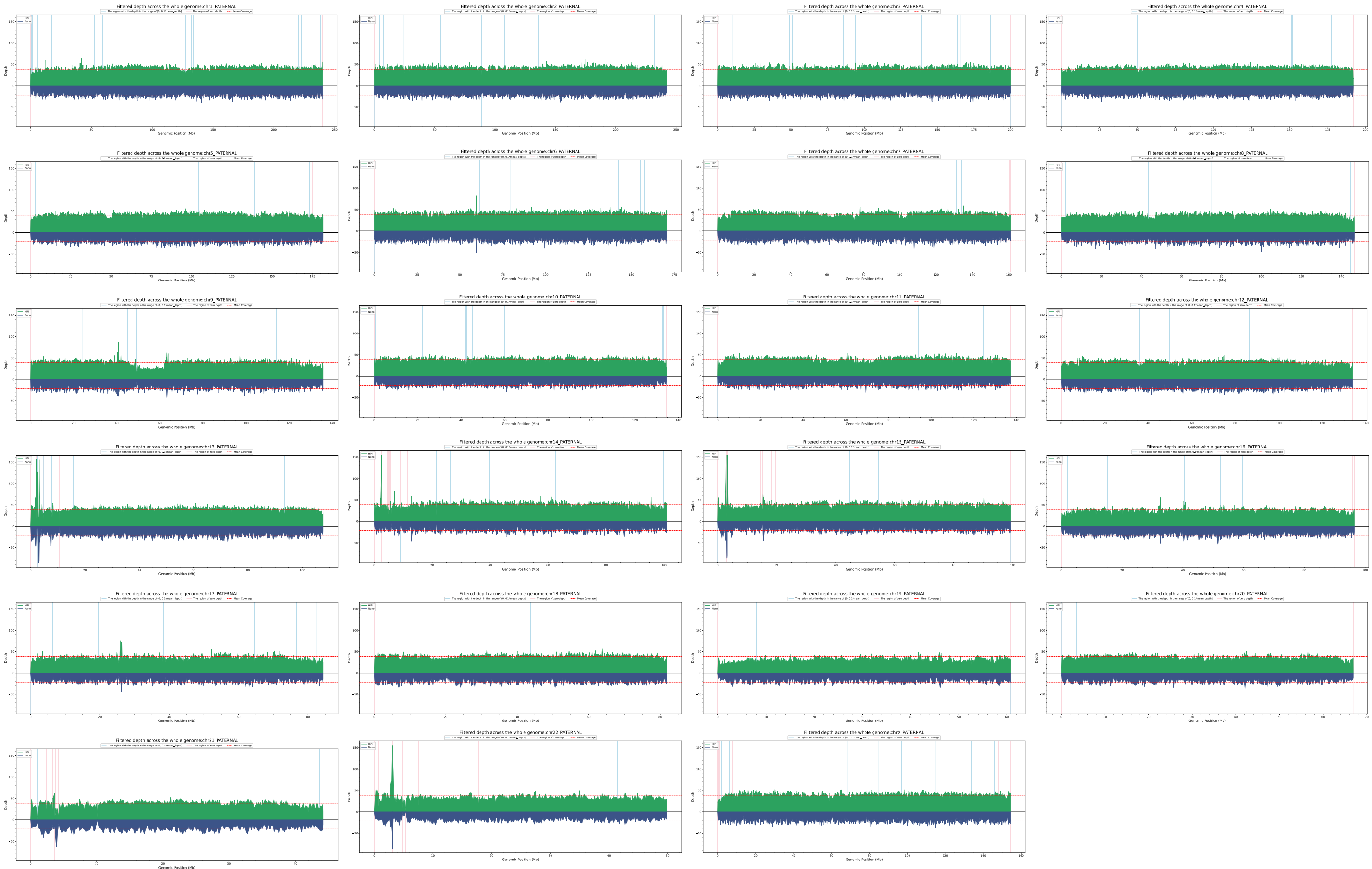


**Supplementary Fig. 15 GCI plot of binned ONT and HiFi reads along the C35.Pat's genome.** GCI, genome continuity inspector.

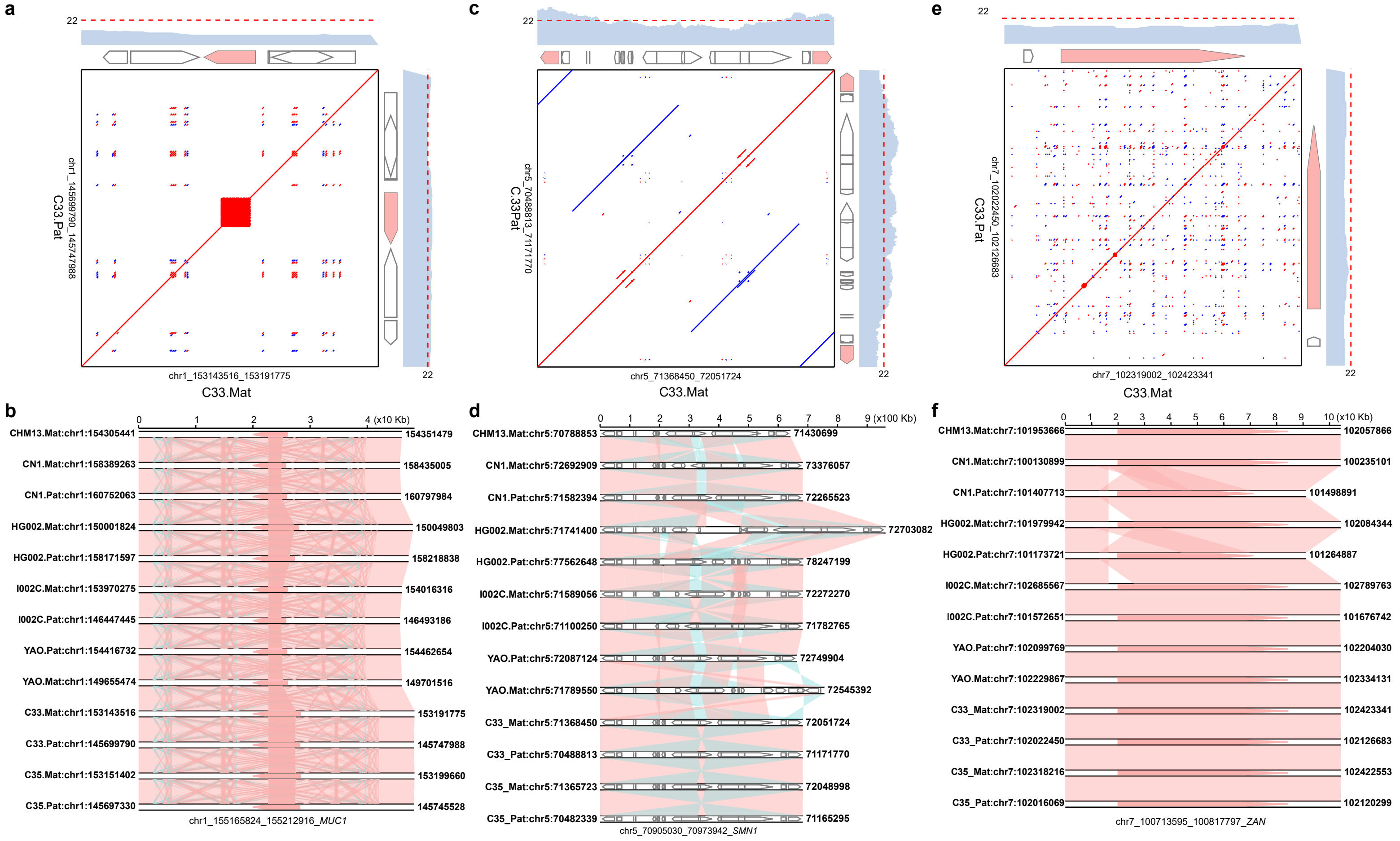


**Supplementary Fig. 16 Sequence coverage of challenging medically-relevant autosomal genes and collinearity of genes in homologous haplotypes. a** Sequence coverage of *MUC1* in homologous haplotypes of twins. **b** Collinearity analysis of *MUC1* in different haplotypes. **c** Sequence coverage of *SMN1* in homologous haplotypes of twins. **d** Collinearity of *SMN1* analysis in different haplotypes. **e** Sequence coverage of *ZAN* in homologous haplotypes of twins. **f** Collinearity analysis of *ZAN* in different haplotypes.

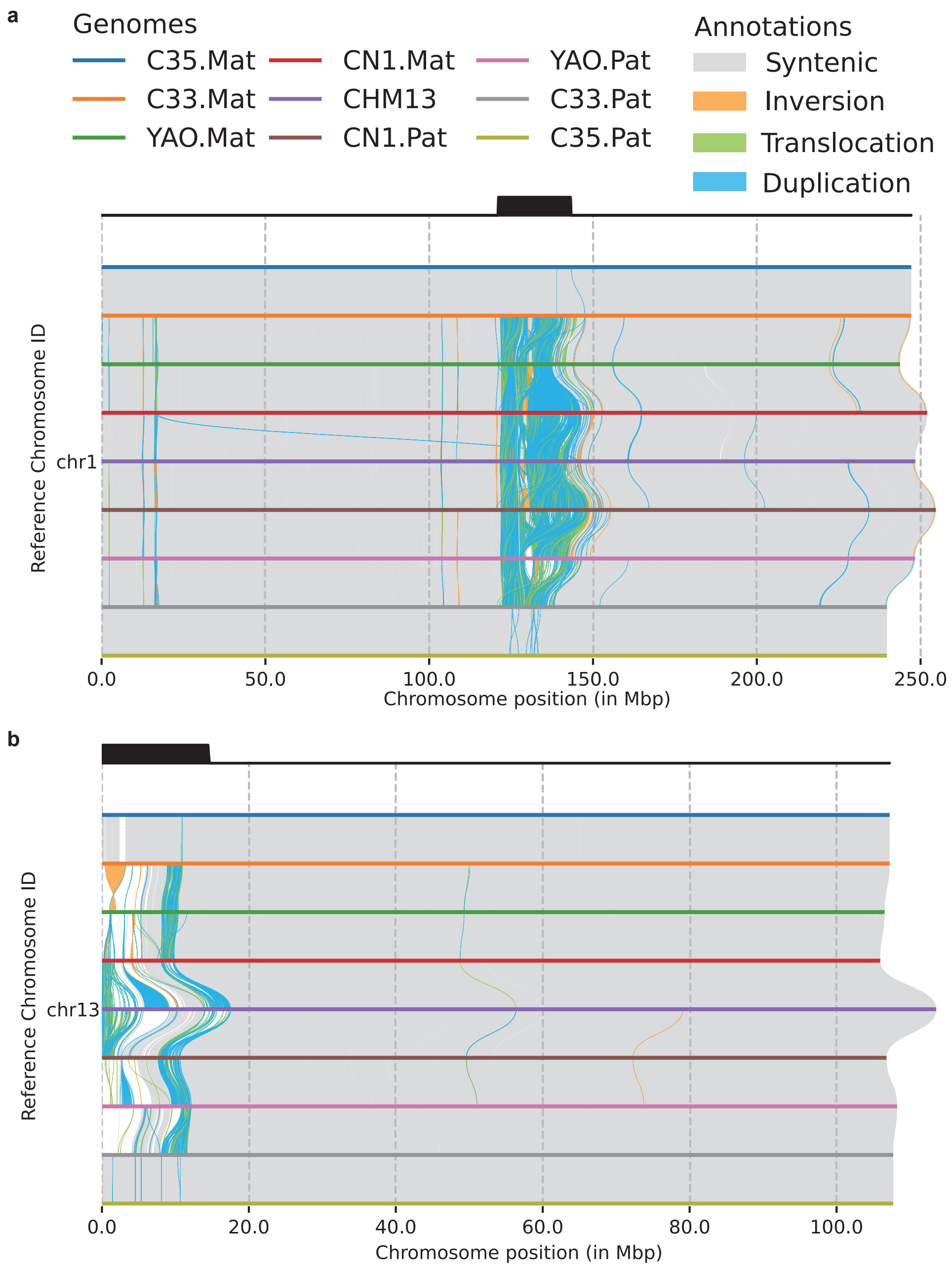


**Supplementary Fig. 17 Collinearity analysis between the twin genome and other reference genomes.** Collinearity analysis of chr1 (a) and chr13 (b) between different haplotypes. The gray lines showed the syntenic regions between haplotypes. The orange, green, and bluish-green lines represented structural variations, specifically inversion, translocation, and duplication, respectively.

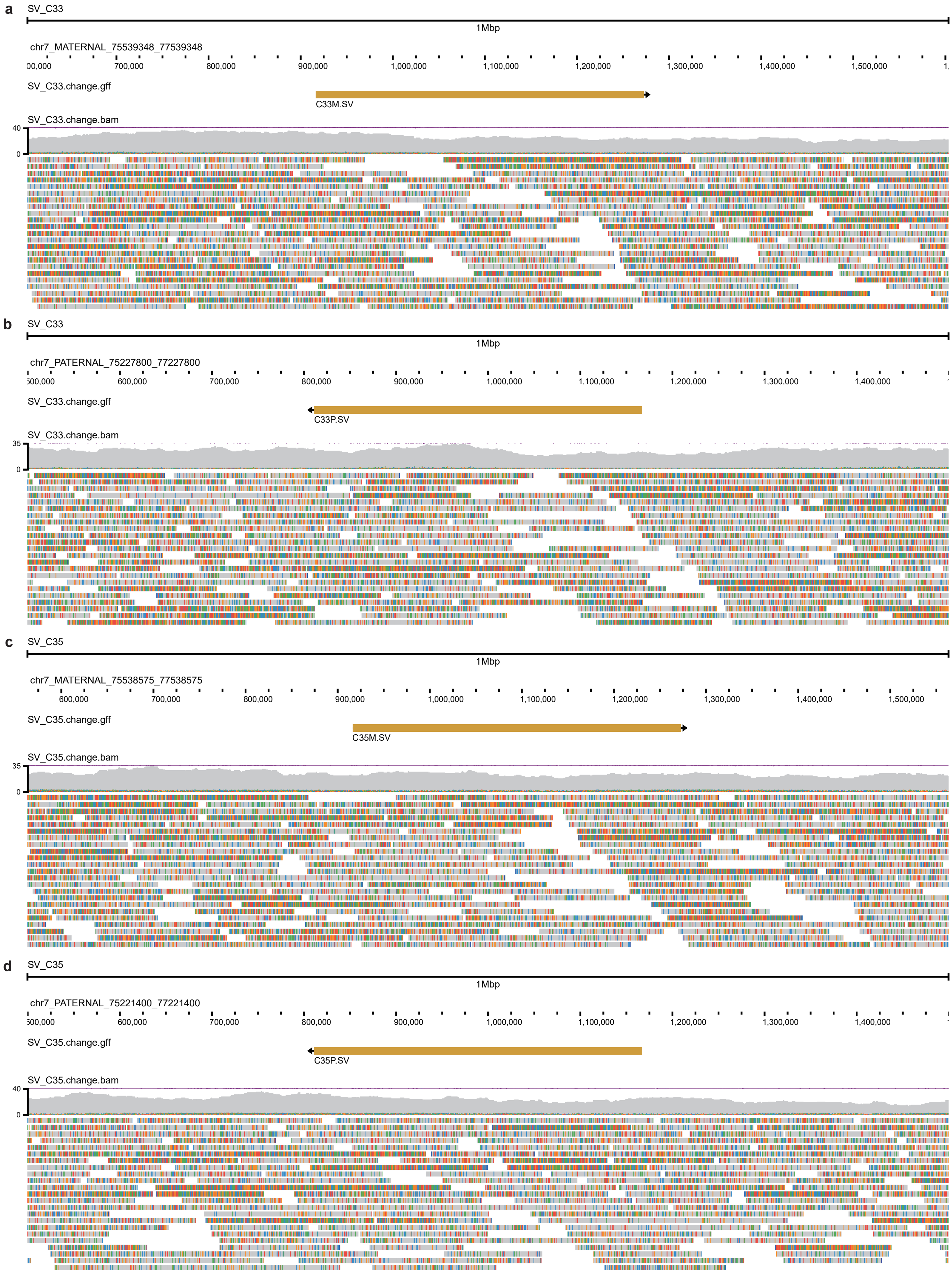


**Supplementary Fig. 18 Comparative IGV visualizations validating genomic assembly accuracy for large-scale inversions on chr7 in C33.Mat (a), C33.Pat (b), C35.Mat (c) and C35.Pat (d).** IGV, Integrative Genomics Viewer.

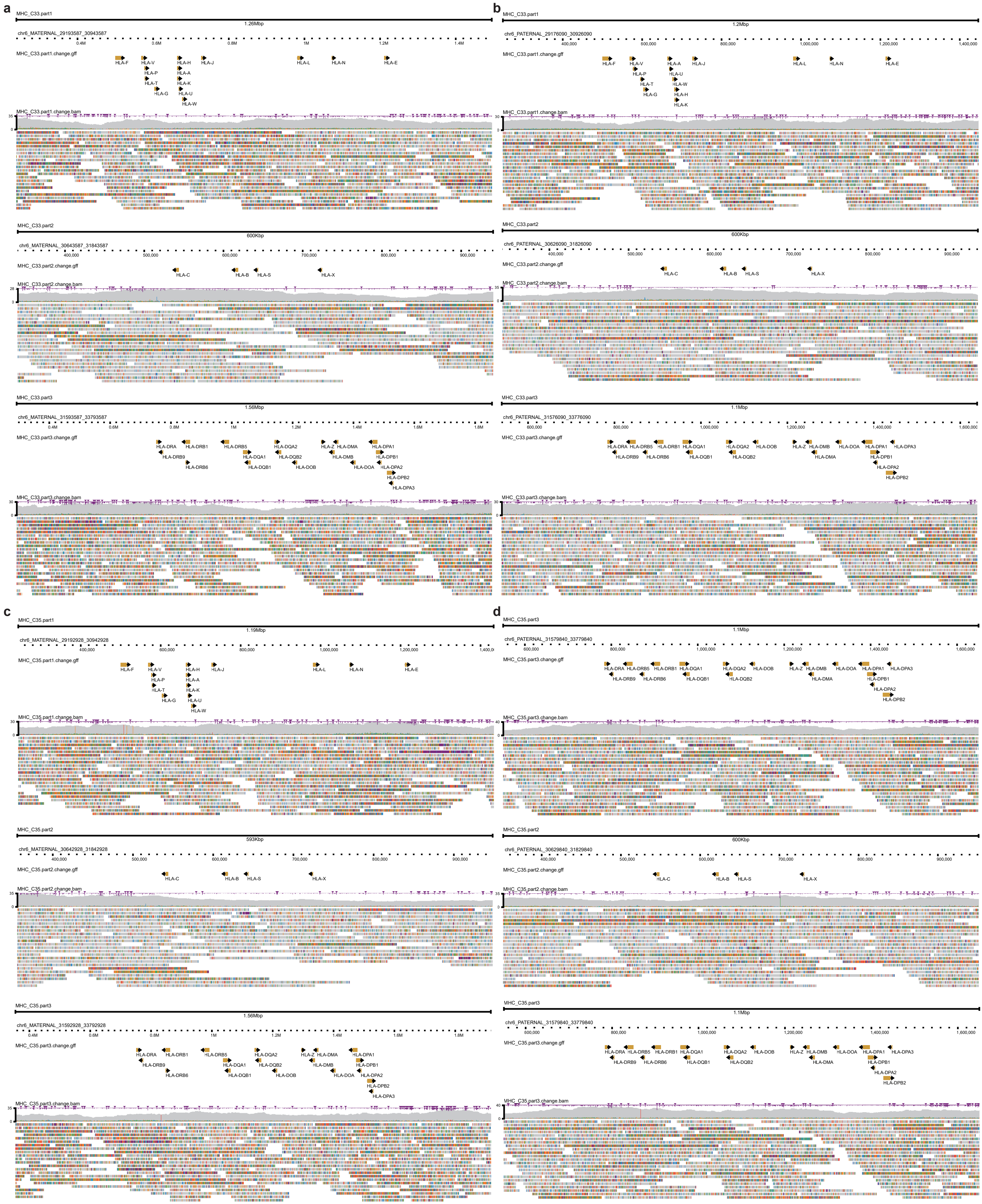


**Supplementary Fig. 19 Comparative IGV visualizations validating genomic assembly accuracy for HLA region in C33.Mat (a), C33.Pat (b), C35.Mat (c) and C35.Pat (d).**

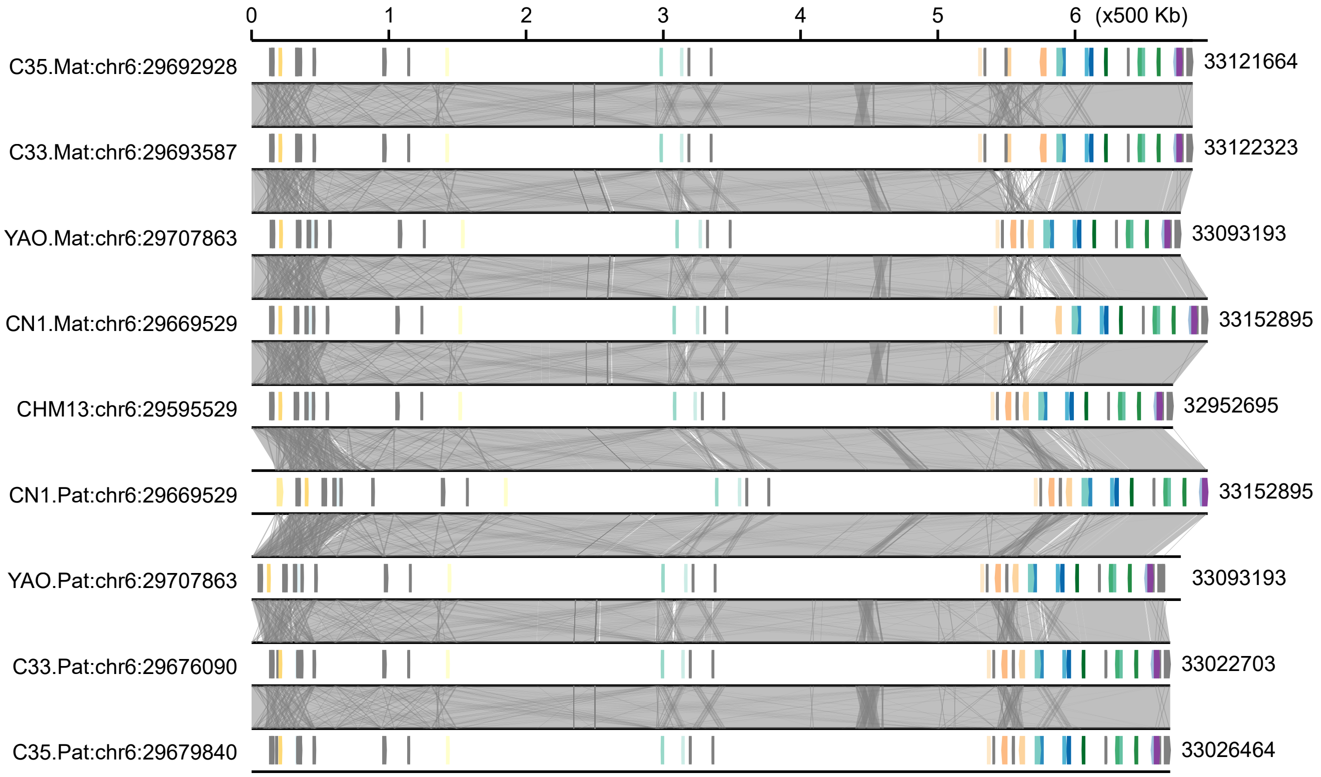


**Supplementary Fig. 20 Collinearity analysis across the genome incorporates HLA gene allelic variations among CN1, YAO, C33, and C35 haplotypes.** Color-coded bars displayed gene annotations for the HLA locus, with each color representing a distinct HLA allele. Gray lines illustrated pairwise collinearity analyses, adn connecting lines highlighted orthologous syntenic blocks.

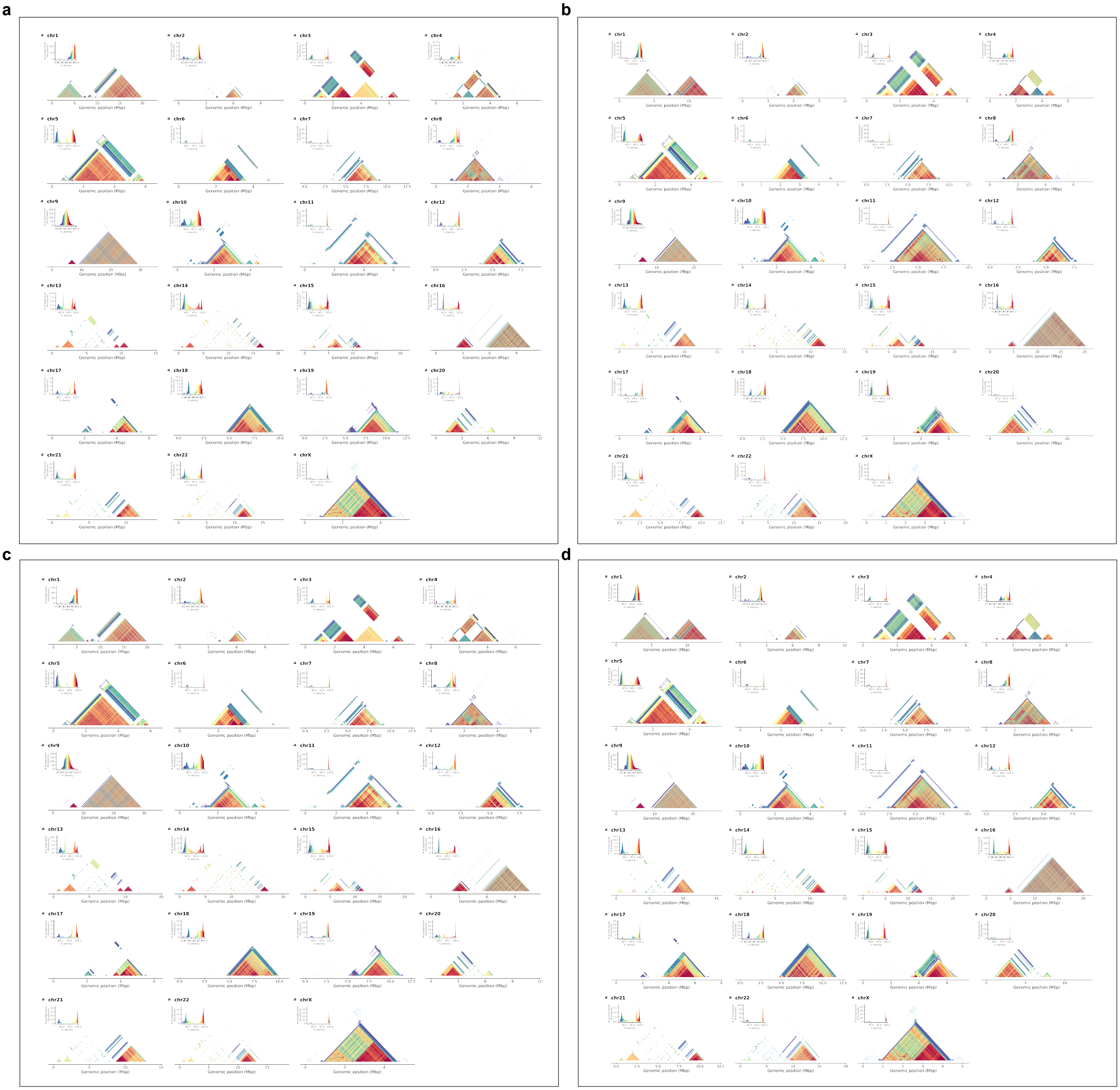


**Supplementary Fig. 21 StainedGlass plots of the CEN regions of C33 and C35.** StainedGlass plots of the CEN regions of chromosomes 1 to 22 and chromosome X in C33.Mat (a), C33.Pat (b), C35.Mat (c) and C35.Pat (d).

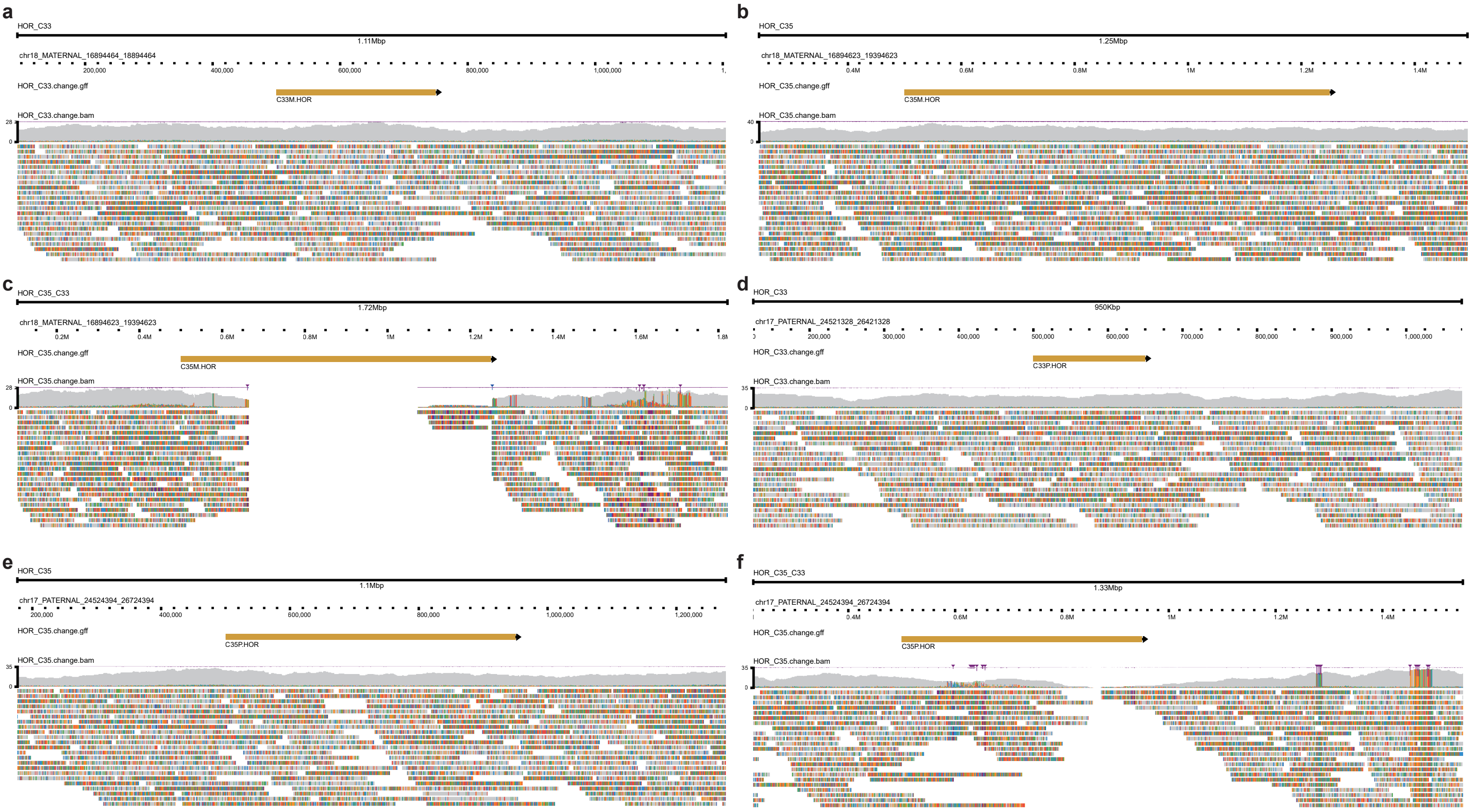


**Supplementary Fig. 22 Comparative IGV visualizations validating genomic assembly accuracy for CNVs in the HOR.** **a** IGV visualization of chr18 in C33.Mat using C33' ONT sequencing data. **b** IGV visualization of chr18 of C35.Mat using C35 ONT sequencing data. **c** IGV visualization generated by mapping ONT reads from C33.Mat to C35.Mat assembly. **d** IGV visualization of chr17 of C33.Pat using C33 ONT sequencing data. **e** IGV visualization of chr17 of C35.Pat using C35 ONT sequencing data. **f** IGV visualization generated by mapping ONT reads from C33.Pat to C35.Pat assembly. IGV, Integrative Genomics Viewer.

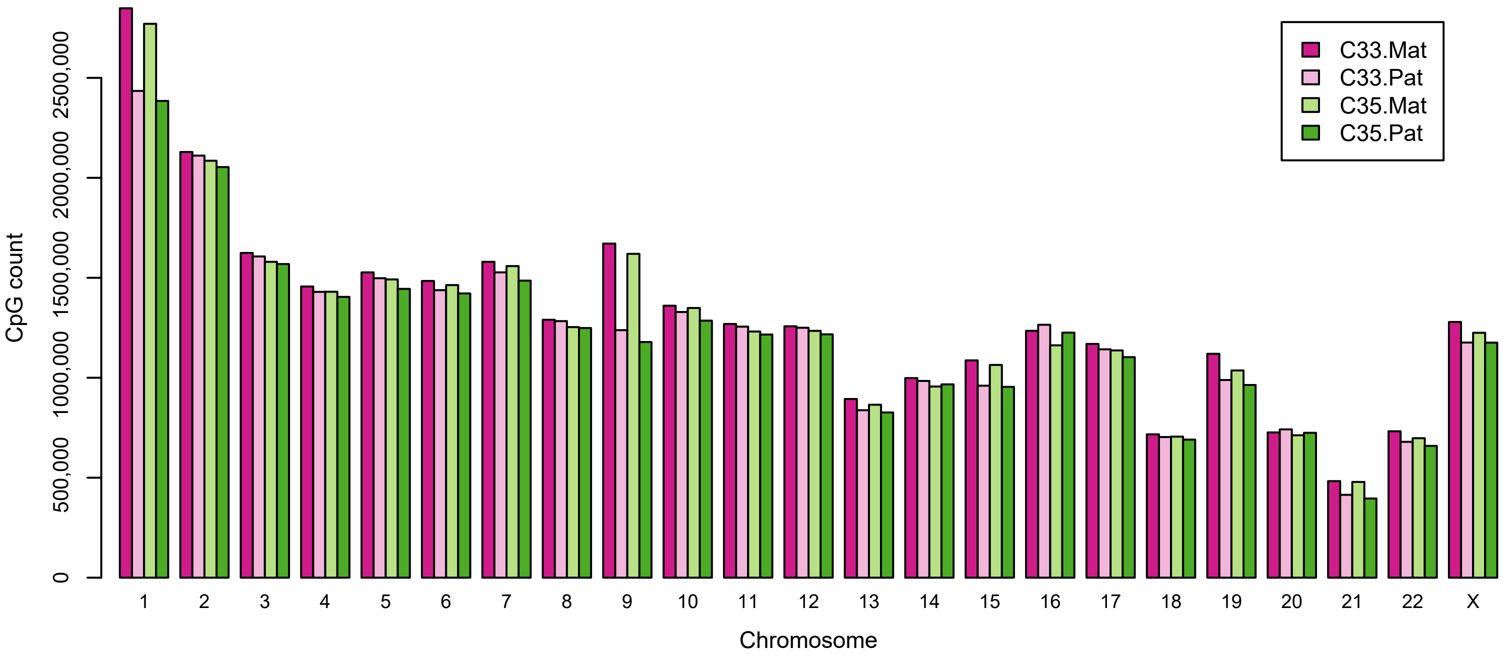


**Supplementary Fig. 23 A bar graph representing the count of CpG sites per chromosome for C33 and C35.**

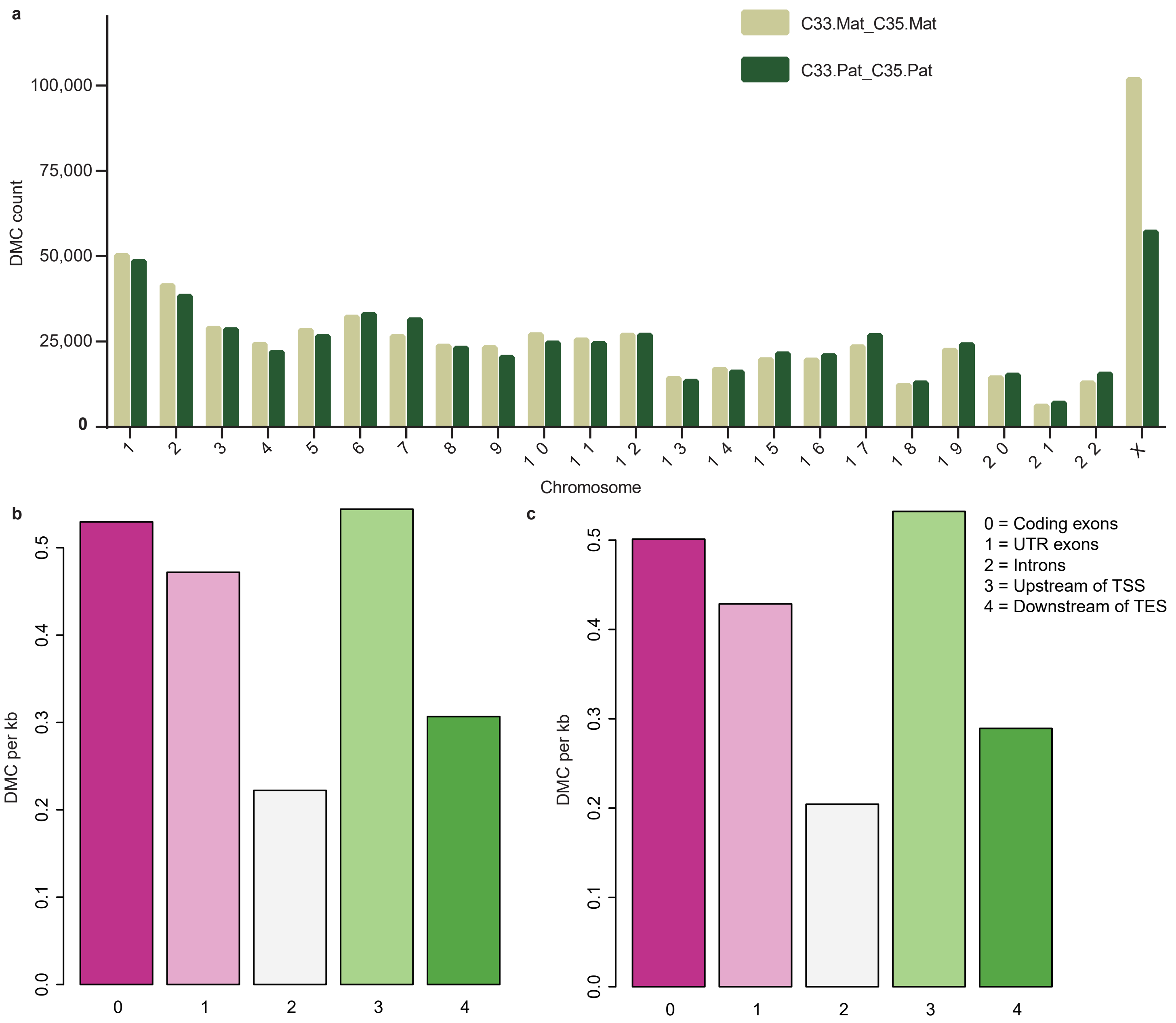


**Supplementary Fig. 24 Genome-wide distribution of DMCs. a** The total counts of DMCs were identified on each chromosome for the maternal (C33.Mat vs. C35.Mat, light yellow) and paternal (C33.Pat vs. C35.Pat, dark green) comparisons. **b** The distribution of DMCs for the maternal haplotypes of twins. **c** The distribution of DMCs for the paternal haplotypes of twins. UTR, untranslated regions. TSS, upstream of transcription start sites. TES, downstream of transcription end sites.

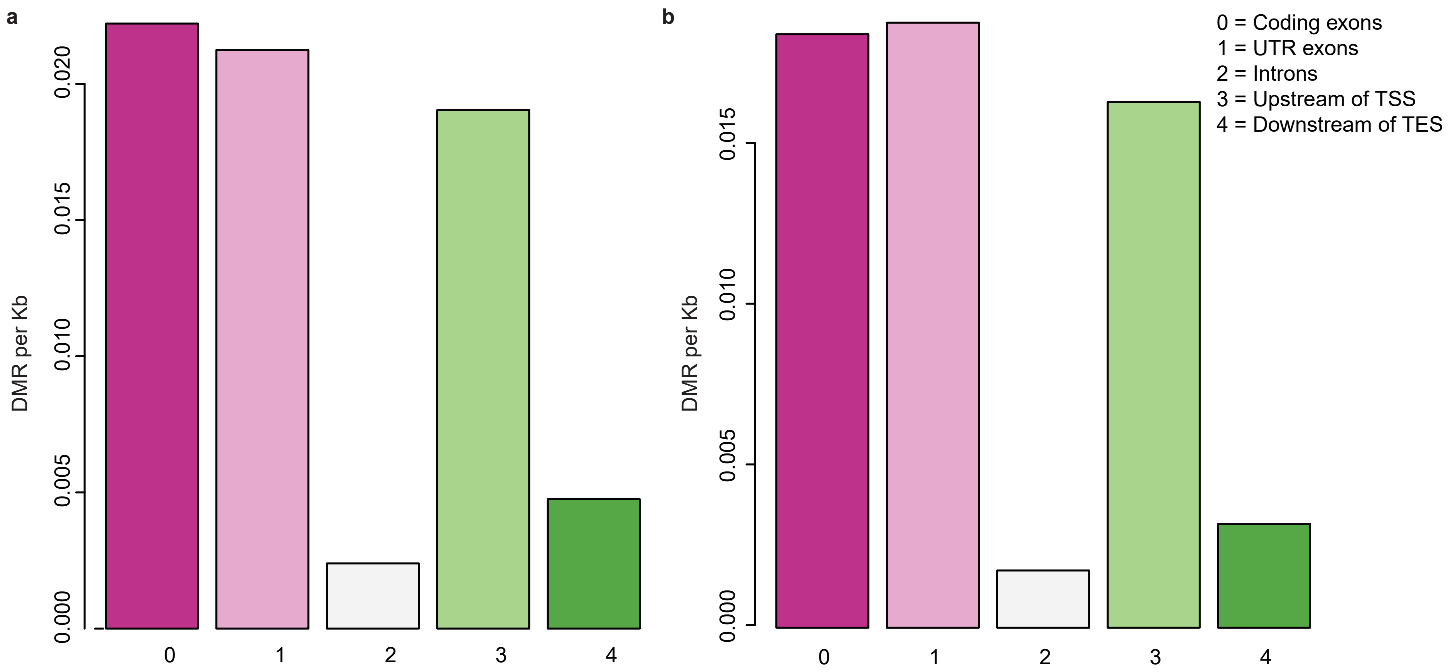


**Supplementary Fig. 25 Genomic distribution of DMRs. a** The distribution of DMRs for the maternal haplotypes of twins. **b** The distribution of DMRs for the paternal haplotypes of twins. UTR, untranslated regions. TSS, upstream of transcription start sites. TES, downstream of transcription end sites.

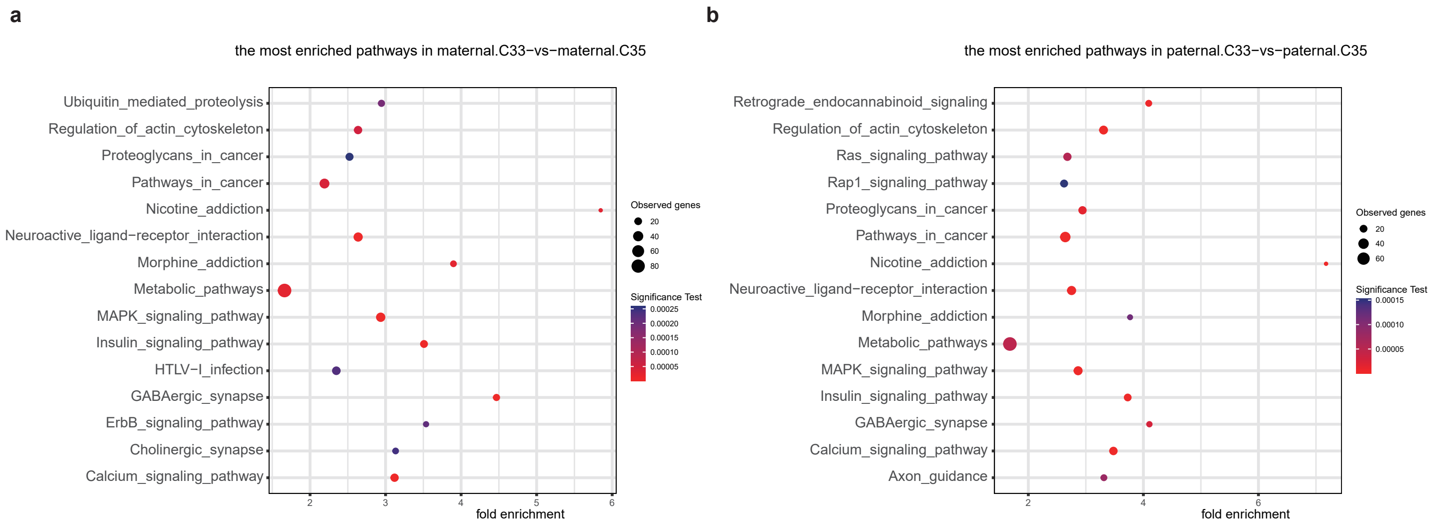


**Supplementary Fig. 26 KEGG enrichment analysis of DMR-associated genes between C33.Mat and C35.Mat (a) and between C33.Pat and C35.Pat (b).**

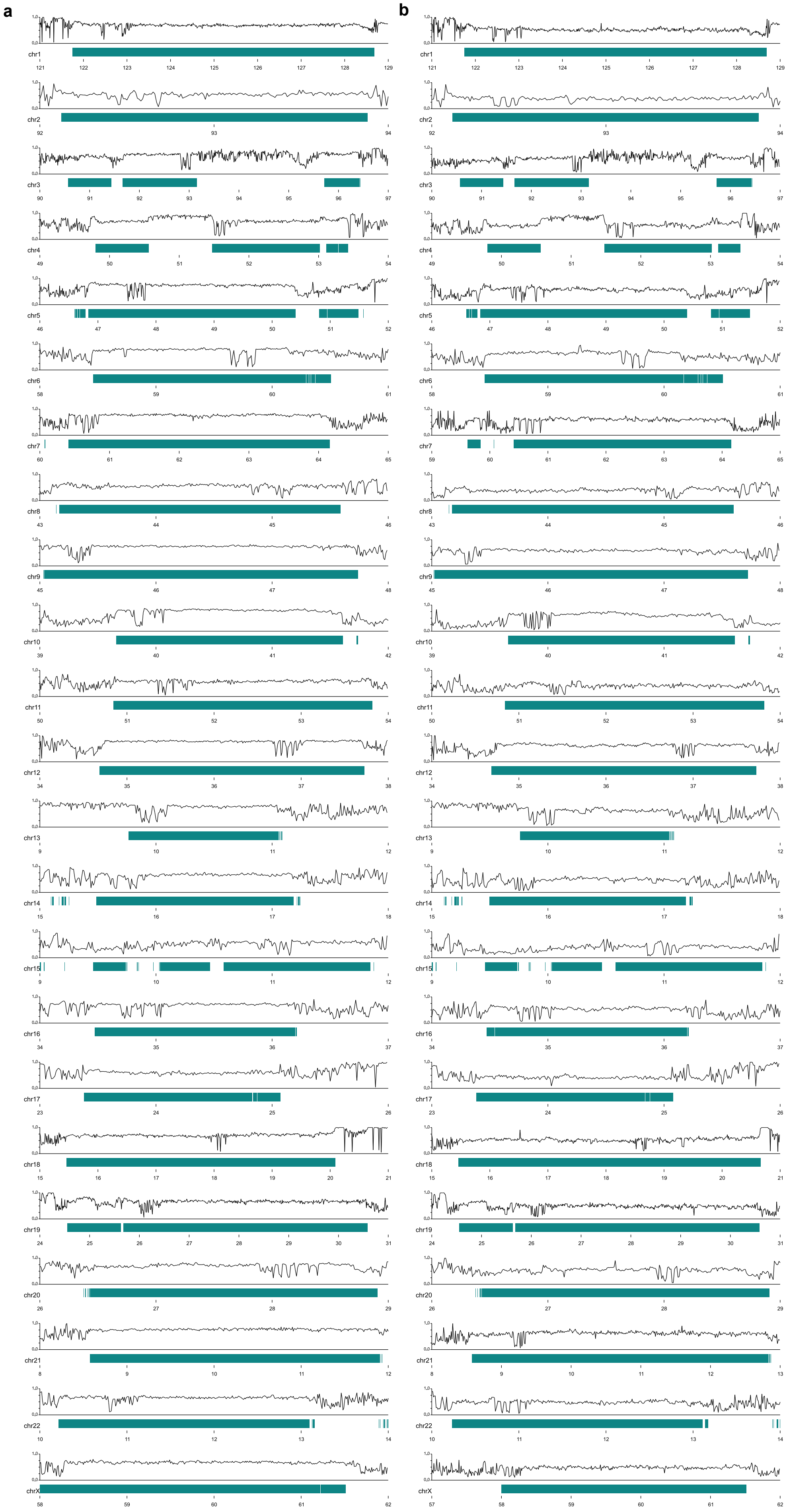


**Supplementary Fig. 27 Methylation levels of C33.Mat (a) and C35.Mat (b) CDR and their flanking region.**

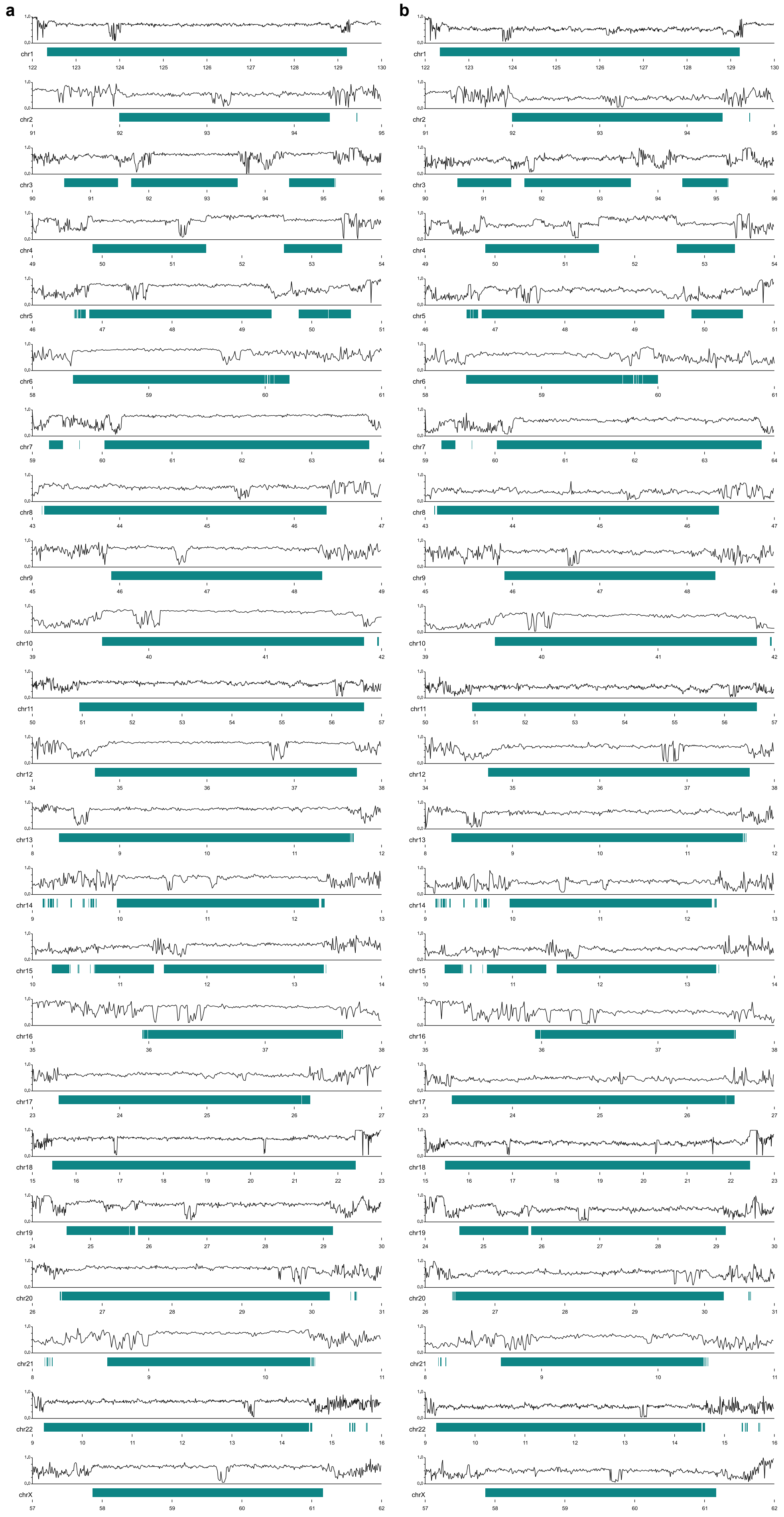

**Supplementary Fig. 28 Methylation levels of C33.Pat (a) and C35.Pat (b) CDR and their flanking region.**

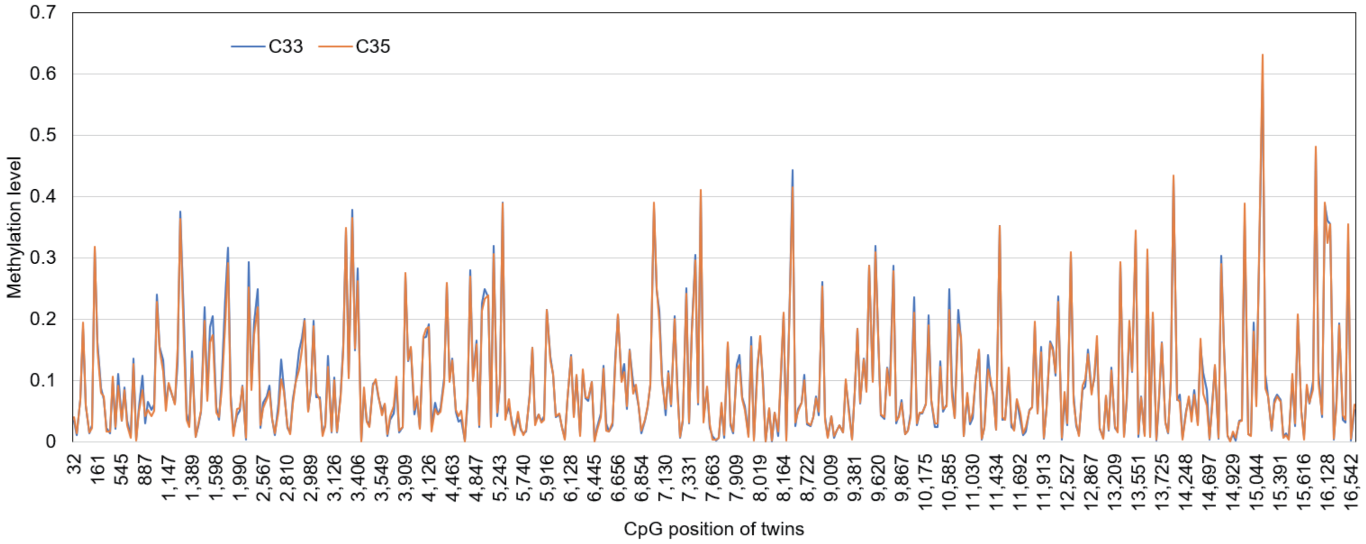


**Supplementary Fig. 29 Methylation profiling of C33 and C35 across the mitochondrial genome.** Chi-square test.

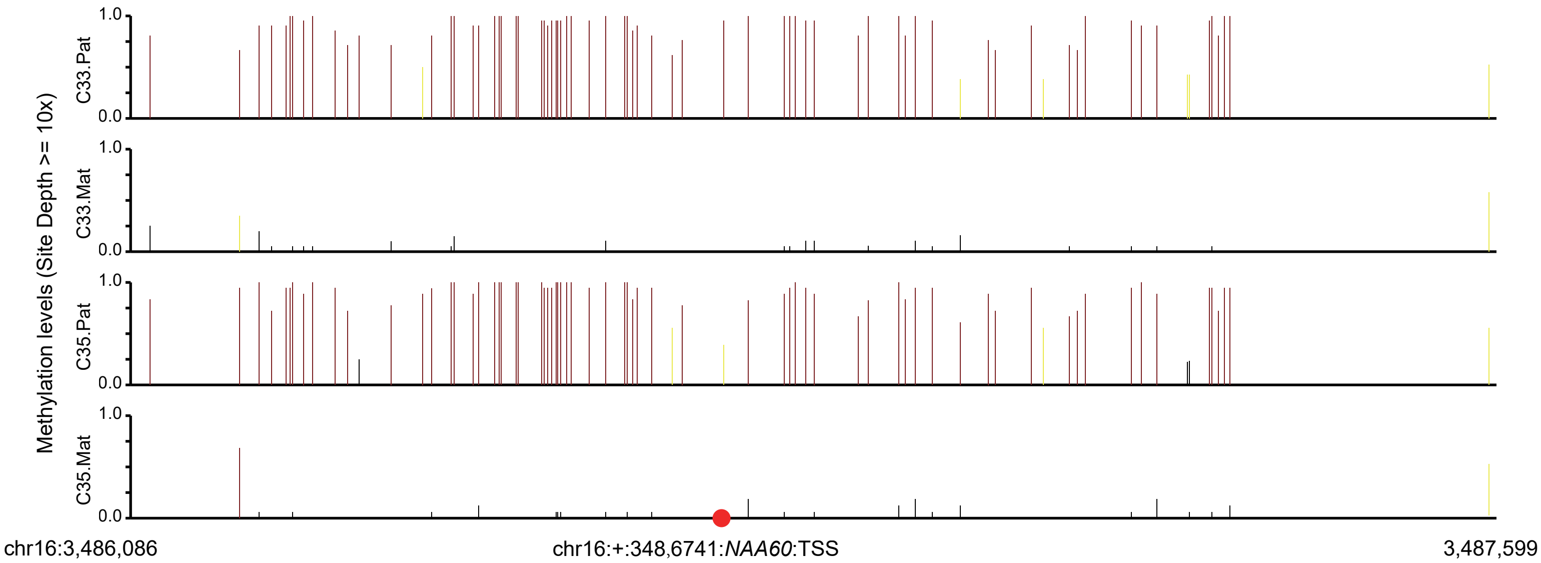


**Supplementary Fig. 30 Differential methylation patterns at NAA60 promoter in monozygotic twins.** ONT sequencing data showed methylation status of individual CpG sites across the NAA60 promoter region on chr16 for the twins. For each individual, data was shown for both the paternal and maternal allele. Each vertical bar represented a single CpG site, with red indicating a methylated site.

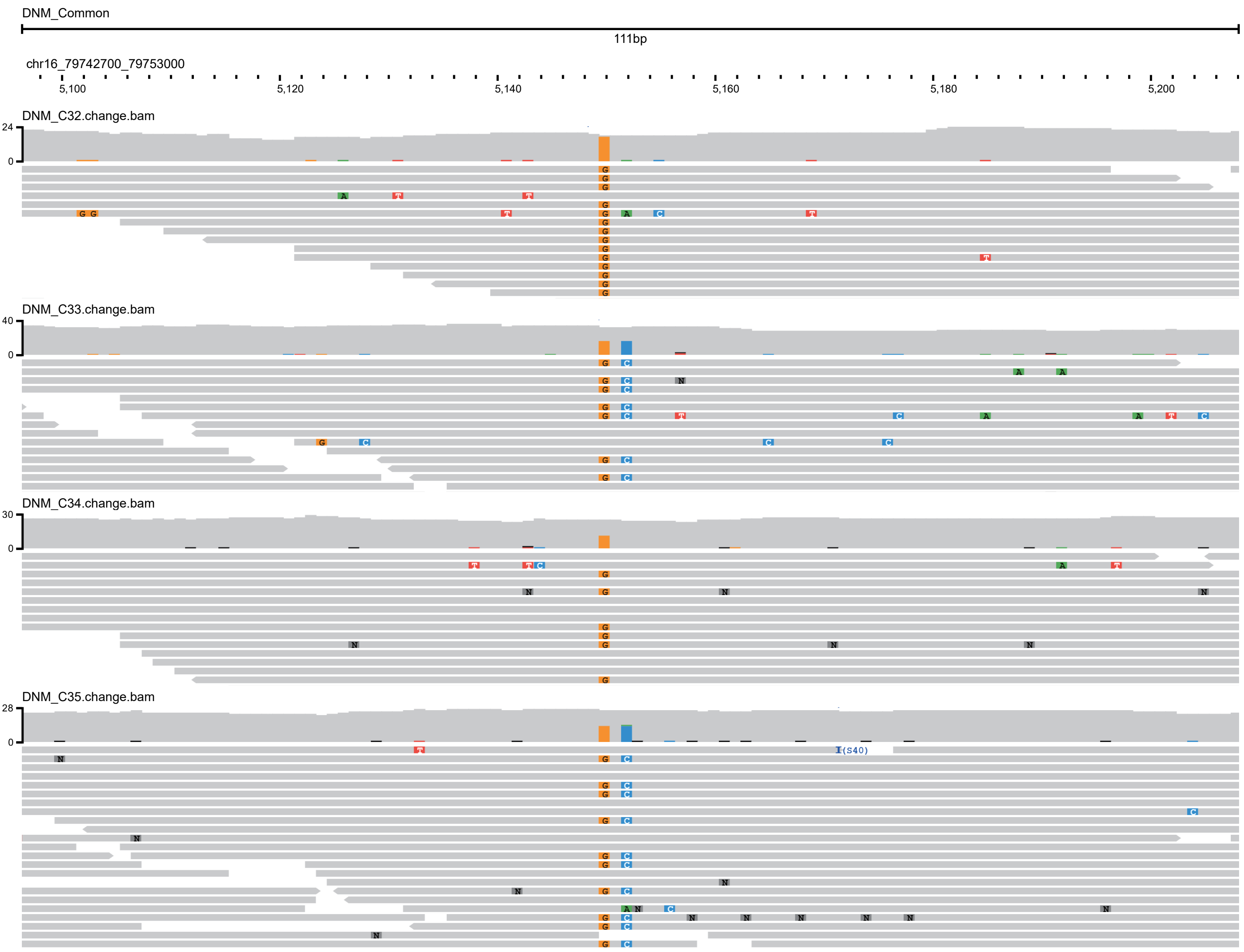


**Supplementary Fig. 31 IGV visualization of *de novo* mutation in *ZFH3* for C33 and C35.**

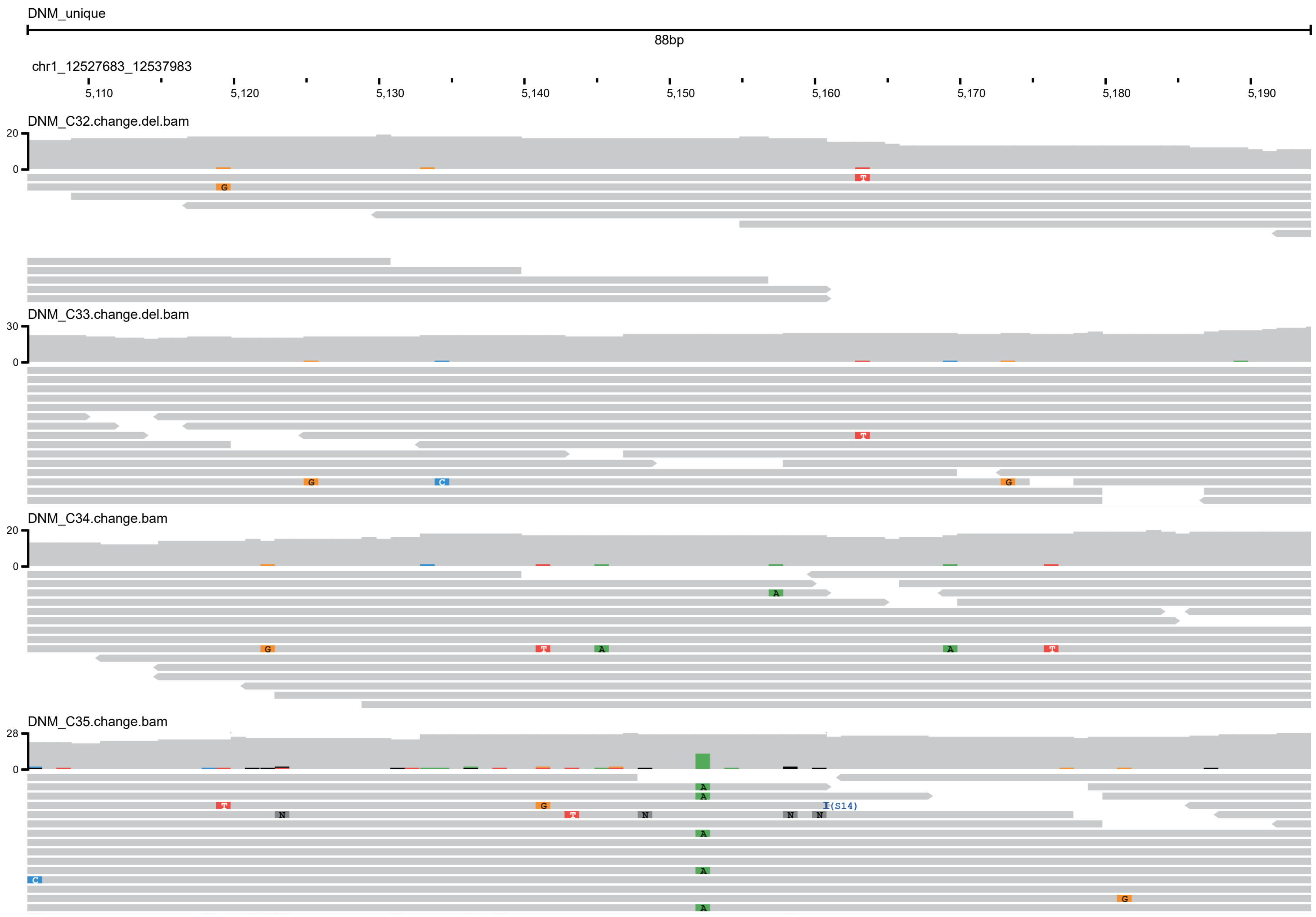


**Supplementary Fig. 32 IGV visualization of de novo mutation in an intergenic region for C33 and C35.**
